# Photoperiod induces sex-specific immune priming in *Pyrrhocoris apterus*

**DOI:** 10.64898/2026.04.27.721165

**Authors:** Adam Bajgar, Gabriela Krejčová, Vlastimil Smýkal, David Doležel

**Affiliations:** Department of Molecular Biology and Genetics, Faculty of Science, University of South Bohemia, 370 05 České Budějovice, Czech Republic; Institute of Entomology, Biology Centre of the Czech Academy of Sciences, 370 05 České Budějovice, Czech Republic

**Author notes:** Corresponding author E-mail addresses (A. Bajgar). Corresponding author (D. Dolezel). These authors contributed equally to this work (Adam Bajgar, Gabriela Krejčová).

**Keywords:** Photoperiodic timer, Immunity, seasonality, immune priming, *Pyrrhocoris apterus*, Insect, hemocytes

## Abstract

Seasonal variation in day length provides a reliable cue that allows insects to anticipate upcoming environmental challenges. Here, we demonstrate that photoperiod induces pronounced, sex-specific immune priming in the linden bug *Pyrrhocoris apterus*. Females exposed to short-day, diapause-inducing conditions exhibited broadly enhanced immune activity compared with long-day females, whereas immune parameters in males were largely unaffected by photoperiod. Short-day females showed increased immune cell abundance, elevated expression of immune-related genes, enhanced humoral immune activity, and increased resistance to bacterial infection. Importantly, photoperiod-induced immune priming depended on a functional *m-cryptochrome* gene, linking seasonal immune regulation to the photoperiodic timer. Consistent with laboratory results, females collected under natural short-day conditions also displayed enhanced immune parameters despite increased environmental variability. Together, our findings identify photoperiod as a key regulator of immune preparedness in female insects and reveal a sex-specific anticipatory immune strategy associated with seasonal timing.

## 1. Introduction

To survive adverse conditions, insects evolved diapause, a genetically programmed arrest of development, metabolism, or reproduction occurring at a species-specific life stage (Schebeck et al., 2024; Kostal., 2006). Prior to diapause, insects build up substantial energy stores, while the ensuing metabolic depression reduces expenditure and facilitates long-term preservation of these reserves. In many insects, seasonal timing is governed primarily by photoperiod, the most reliable environmental signal of impending change. Light information perceived by photoreceptors is processed by a photoperiodic timer (also known as photoperiodic clock), which measures day length and regulates the decision to enter or avert diapause (Kostal, 2011; Denlinger et al., 2017; Saunders, 2020; Doležel, 2015).

Diapause is accompanied by broad physiological remodeling, including changes in reproductive organs, digestive function, fat body organization, and the accumulation of storage and protective compounds in the hemolymph (Denlinger et al. 2002). These adjustments enhance survival during prolonged exposure to low temperature, food limitation, and other winter-associated stresses. Because seasonal adaptation also alters behavior and likely changes pathogen exposure, it is reasonable to expect that immunity is integrated into the photoperiodic response (Kostal, 2006).

Activation of the insect immune system is commonly associated with changes in the abundance, mobilization, and differentiation of circulating hemocytes into functionally specialized cell types (Lavine and Strand 2002). This cellular response is accompanied by activation of humoral immunity, including increased production of antimicrobial peptides and the induction of enzymatic pathways involved in melanization and hemolymph coagulation (Eleftherianos et al. 2021). Although insect immunity lacks the adaptive branch characteristic of vertebrates, it nevertheless provides highly effective protection against infection. In addition, it is now well established that prior pathogen exposure can enhance protection upon subsequent challenge, a phenomenon referred to as immune priming (Krejcova and Bajgar 2025). Depending on the system, this enhanced responsiveness may involve changes in hemocyte number or activity as well as altered expression of humoral immune factors (Sulek et al. 2021). By analogy, seasonal adjustment of immunity toward a pre-activated or preconditioned state may represent an important strategy for improving survival during predictable periods of elevated stress.

A number of studies suggest that seasonal state is associated with changes in insect immunity, but the available evidence remains fragmented and often difficult to interpret mechanistically. Early hemocytological studies showed that diapause may be accompanied by substantial reorganization of the cellular immune compartment. In *Pectinophora gossypiella*, diapausing larvae displayed reduced total and differential hemocyte counts, which increased again after diapause termination (Raina and Bell, 1974), and similar reductions were reported in *Sesamia cretica* (El-Mandarawy, 1997). However, diapause does not necessarily imply a general shutdown of immune competence, as diapausing pupae of *Samia cynthia pryeri* retained hemocyte-mediated phagocytosis and encapsulation (Nakamura et al., 2011). Seasonal immune responses also appear highly species specific: in overwintering larvae, Ferguson and Sinclair (2017) found stable hemocyte numbers in *Curculio* sp. and *Eurosta solidaginis*, but reduced hemocyte abundance and antimicrobial activity in *Pyrrharctia isabella*. Interpretation is further complicated by developmental stage, because in larvae and pupae hemocytes contribute not only to immunity but also to growth, tissue remodeling, and metamorphosis (Krejcova et al. 2024). Importantly, most of these studies do not isolate photoperiod as an independent causal factor, as they compare diapausing and non-diapausing individuals, overwintering stages, or field-collected insects sampled across seasons. Thus, while seasonal immune remodeling is increasingly well documented, the specific contribution of photoperiod per se remains insufficiently resolved. Consistent with a direct role of day length, Lindberg et al. (2025) showed that *Acheta domesticus* adults maintained under defined light:dark regimes differed in several immune-associated hemolymph parameters, although such direct experimental evidence remains scarce.

An excellent model to address this question is the linden bug, *Pyrrhocoris apterus*, which has been studied extensively as a model of both circadian and photoperiodic timing. In this species, reproductive and diapause phenotypes can be induced under constant thermal conditions solely by changing day length, making it possible to isolate the effect of photoperiod from other environmental variables (Socha et al. 2013; Saunders, 1983). Under long-day conditions (LD, 18 h light : 6 h dark), adults develop a reproductive phenotype, whereas under short-day conditions (SD, 12 h light : 12 h dark), they enter reproductive diapause (Hodkova and Hodek 1994). These alternative seasonal states differ in multiple physiological traits, including the status of the reproductive organs, digestive tract, fat body, and hemolymph composition (Smykal et al. 2014; Bajgar et al., 2013a,b). components of the melanization pathway Photoperiodic signaling in *P. apterus* depends on components of the circadian system, and a functional *mammalian-type cryptochrome* (*m-cry*) is required for proper photoperiodic responses, whereas the *Drosophila-type cryptochrome* is absent in this species and the entire infraorder Pentatomomorpha (Kotwica-Rolinska et al. 2022; Bullo et al., 2025).

Downstream translation of photoperiodic information into seasonal physiology is strongly influenced by endocrine signaling, with juvenile hormone playing a central role and *corpora allata* activity representing an important regulatory node (Hodkova and Okuda 2019, Bajgar et al. 2013a). At the same time, available evidence suggests that diapause entry and termination are regulated differently in males and females, indicating sexually dimorphic downstream mechanisms, although their precise nature remains unresolved (Hejnikova et al. 2022). Despite the extensive use of *P. apterus* as a model for photoperiodism and diapause, relatively little is known about its immune system. Current knowledge is largely limited to the identification of several inducible antimicrobial factors and a basic classification of circulating hemocytes (Berger and Jurcova 2012, Berger and Slavickova 2008). Whether and how photoperiod shapes immune organization and function in this species has not been systematically addressed.

In this study, we investigated how photoperiod shapes immune system organization and function in *P. apterus* by combining controlled laboratory experiments, genetic disruption of photoperiod perception, and analyses of field-collected individuals. We show that SD conditions induce a pronounced enhancement of immune preparedness in females, but not in males, and that this response depends on a functional mammalian-type cryptochrome. Our results identify photoperiod as a strong regulator of both cellular and humoral immunity in *P. apterus* and reveal a sex-specific anticipatory immune strategy associated with seasonal timing.

## 2. Materials and methods

### 2.1. Insect rearing, strains, and photoperiodic regimes

Laboratory-reared wild-type linden bugs of Oldrichovec strain (Pivarciova et al., 2016) and CRISPR/Cas9-induced *m-cry*^*04*^ mutants (Kotwica-Rolinska et al., 2019; Kotwica-Rolinska et al., 2022) were maintained in climate-controlled incubators at a constant temperature of 25 °C and relative humidity of 60%. Individuals were assigned to specific photoperiodic regimes starting from the egg stage. The short-day (SD) regime consisted of 12 h light and 12 h dark (12L:12D) and induced reproductive diapause, with adults remaining reproductively inactive. In contrast, insects maintained under long-day (LD) conditions (18 h light and 6 h dark; 18L:6D) were reproductively active. Insects were sacrificed 4-7 hours after light on signal (Zeitgeber time, ZT 4-7). Experimental individuals entered the study as adults (imago) six days after the final molt. Experimental animals were fed linden seeds and had ad libitum access to water. Field-caught individuals were collected in the South Bohemian region (Czech Republic) at two locations: the Stromovka city park (GPS - 48.972201, 14.452195) and the vicinity of the Svět fishpond near Třeboň (GPS - 49.000455, 14.772287). Long-day field-caught (LD-FC) individuals were collected between 21 and 25 June 2022, between 10:00 and 13:00 hour each day (ZT 5-8), when the daily average temperature was about 21°C and local photoperiodic conditions closely matched an 18L:6D regime. SD field-caught (SD-FC) individuals were collected between 22 and 26 September 2022, between 10:00 and 13:00 hour each day (ZT 3-6), when the daily average temperature was 10°C (early morning minima around 4-7°C) and natural photoperiodic conditions approximated a 12L:12D regime. Prior to experimentation, wild-caught insects were maintained under outdoor conditions.

### 2.2. Hemolymph sampling

Hemolymph was collected by cutting the tip of an antenna, followed by gentle pressure applied to the body to extrude a droplet of hemolymph containing circulating hemocytes. The hemolymph droplet was immediately collected into a defined volume of ice-cold phosphate-buffered saline (1xPBS; 137 mM NaCl, 2.7 mM KCl, 10 mM phosphate buffer, pH 7.4) and kept on ice. For each sample hemolymph from six individuals was collected. Samples were subsequently processed according to the requirements of downstream analyses. For microscopic analyses, hemolymph samples were placed directly onto microscope slides for further staining procedures. For gene expression analyses, hemolymph was immediately mixed with an excess of TRIzol reagent and snap-frozen in liquid nitrogen, followed by storage at −80 °C until further processing. When required for assays necessitating the removal of hemocytes, cells were separated from the hemolymph by centrifugation at 10,000 × g for 10 min at 4 °C.

### 2.3. Staining, confocal microscopy, and hemocyte quantification

For confocal microscopy, 10 µl of hemolymph was applied onto a microscope slide within a defined square area (1 × 1 cm). Cells were allowed to adhere to the slide for 20 min to promote cell spreading and facilitate morphological differentiation. To distinguish individual hemocyte types, cells were stained with structural dyes: Toluidine blue (1:1), nuclei were labeled with DAPI (1:10,000), and the actin cytoskeleton was visualized using Phalloidin Red (Thermo Scientific; 1:1,000). To assess the phagocytic activity of individual hemocyte classes, insects were injected with 2 µl of the phagocytic marker *S. aureus* pHrodo™ (Thermo Scientific; 1:1,000) prior to hemolymph isolation. In these experiments, Phalloidin Green (Thermo Scientific; 1:1,000) was used as a cytoskeletal marker. All circulating cells present in the hemolymph samples were examined for their characteristic morphological features and classified into hemocyte subpopulations based on cytoskeletal organization, nuclear size, and phagocytic activity. The classification of individual hemocyte subpopulations was based on previous studies (Berger and Jurčová, 2012; Cociancich et al., 1994). Confocal imaging was performed using an Olympus FluoView 3000 confocal microscope to obtain high-resolution images and detailed morphological information. Quantification of hemocytes was conducted using an inverted fluorescence microscope (Olympus BX63). For each defined square area, ten non-overlapping regions of interest were randomly selected, and hemocytes were counted manually. Cell numbers were normalized to 1 µl of collected hemolymph. Confocal images were processed using FIJI software.

### 2.4. Gene identification and primer design

To identify candidate components of the *P. apterus* immune response, we used the same approach as in our recent search for circadian clock genes in Heteroptera (Smýkal et al., 2025). First, we performed a TBLASTN search of the *P. apterus* transcriptome shotgun assemblies (TSA) using *Drosophila melanogaster* protein sequences as queries, yielding several top hits. Protein sequences encoded by TSAs were annotated using the InterProScan plugin (Quevillon et al., 2005) within Geneious Prime to identify protein domains. Reciprocal BLAST searches were then used to expand the dataset with sequences from other insect representatives. All retrieved sequences were aligned using MAFFT, and FastTree analysis was used to identify *P. apterus* duplicated genes. Unequivocally identified protein sequences of genes of interest were then blasted against in-house *P. apterus* Oxford Nanopore Technology (ONT) transcriptomic databases and draft genomic contigs to reveal isoforms and exons within transcripts. Protein sequences were aligned using the MAFFT E-INS-i algorithm, and phylogenetic trees were inferred using the RAxML algorithm (Stamatakis 2006) (Geneious Prime). See Supplementary material for sequences of one representative transcript per measured immune response gene, including transcripts and a consensus sequence of seven *hemiptericin* paralogs.

Primers for transcript quantification were designed to target an amplicon that is present in all splicing isoforms of the gene. Primers to quantify *hemiptericins* bind regions conserved in all seven paralogs.

### 2.5. RNA isolation, reverse transcription, and quantitative PCR

Total RNA was isolated using TRIzol reagent following a standard RNA extraction protocol. To improve pellet visualization and enhance RNA recovery from relatively small amounts of starting material, 1 µl of RNase-free glycogen (1 µg/µl) was added prior to RNA precipitation with isopropanol. RNA concentration and quality were assessed spectrophotometrically.

Reverse transcription was performed using the SuperScript™ III First-Strand Synthesis System (Invitrogen) with oligo(dT) priming, according to the manufacturer’s instructions.

Quantitative PCR (qPCR) was carried out using a SYBR Green–based detection system (2× SYBR Master Mix, Top-Bio). Genes selected for expression analysis were chosen based on established knowledge of insect immunity and identified in the *P. apterus* transcriptome. Primer pairs were initially tested on pooled cDNA samples, and only those producing a single amplicon of the expected size and exhibiting adequate amplification efficiency were used for further analyses.

Gene expression levels were normalized to the ribosomal protein gene *RP49*, which has been routinely used in our laboratory as a stable housekeeping gene.

**Table 1.**
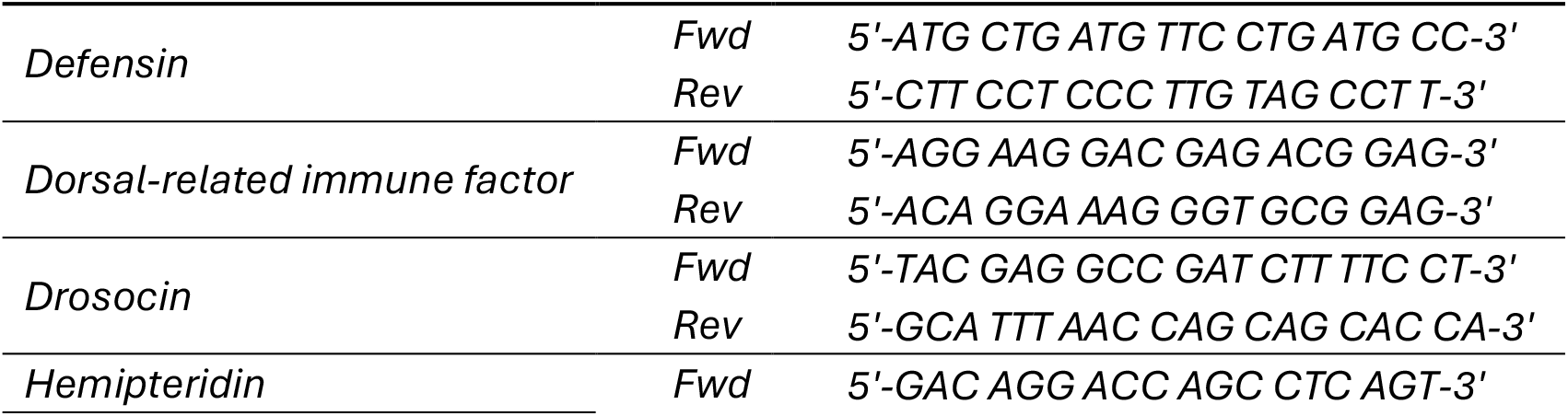
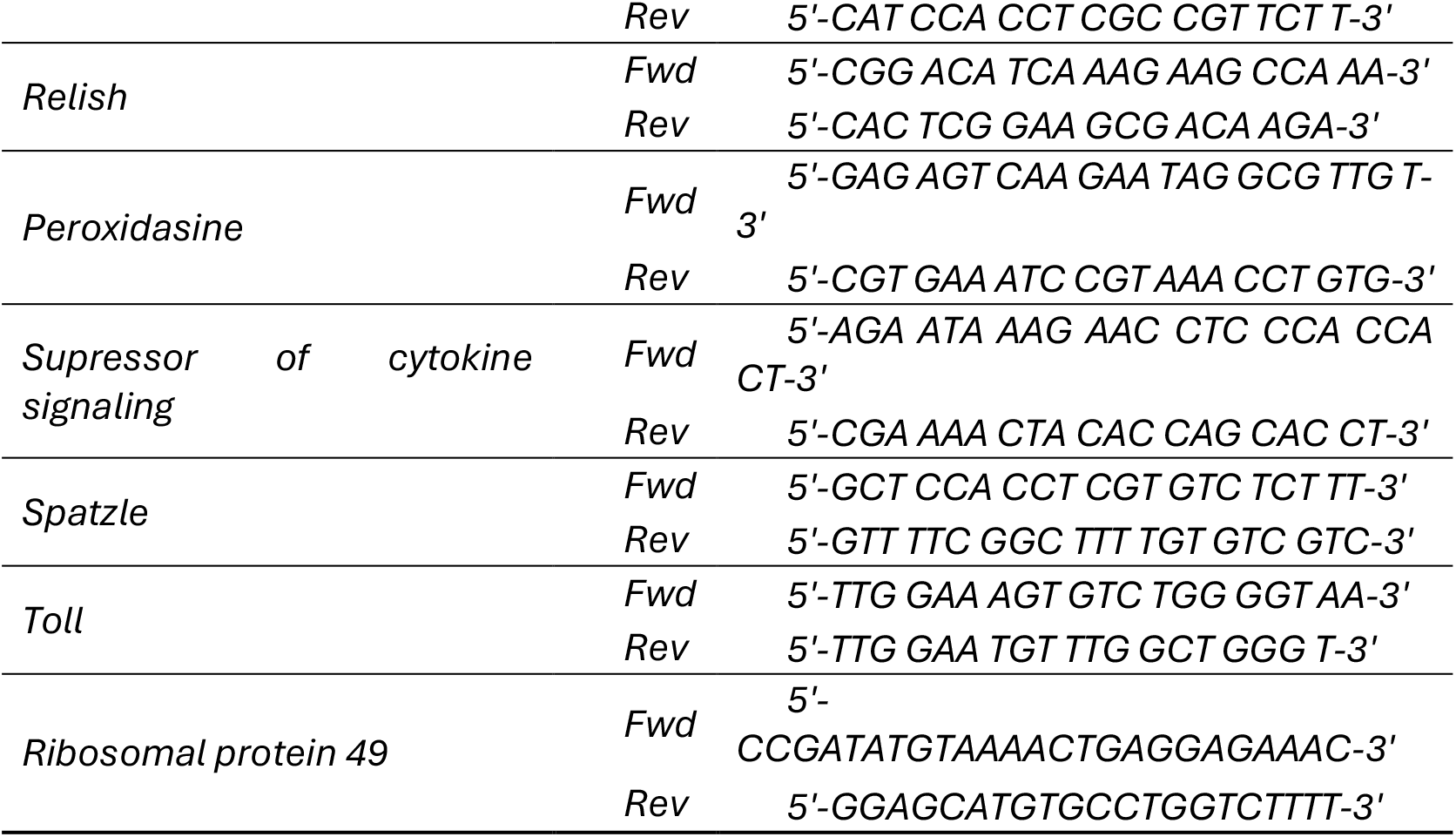
List of primers used for qPCR analysis.

### 2.6. Phenoloxidase assay

Phenoloxidase activity in hemolymph was measured using an L-DOPA–based spectrophotometric assay (Sorrentino et al. 2002). Hemolymph (10 µl) was collected from individual insects and immediately diluted in 160 µl of 10 mM sodium phosphate buffer. For each reaction, 20 µl of diluted hemolymph was combined with 20 µl of L-DOPA (Sigma; 2 mg/ml in 10 mM sodium phosphate buffer) and 60 µl of additional sodium phosphate buffer in a 96-well plate, resulting in a final reaction volume of 100 µl. The formation of dopachrome was monitored as an increase in absorbance at 495 nm using a SunRise microplate reader (Tecan). Absorbance was recorded kinetically with 20 consecutive measurements at 3-min intervals. Based on time-course analyses, absorbance values obtained approximately 80 min after reaction initiation were used for comparative analyses, as this time point provided the highest discriminatory power between samples. Data are presented as relative absorbance values to enable comparison of phenoloxidase activity among samples.

### 2.7. Cultivation of Streptococcus pneumoniae

The *Streptococcus pneumoniae* strain EJ1 (a streptomycin-resistant derivative of D39) was stored at −80 °C in Tryptic Soy Broth (TSB) supplemented with 10% glycerol. For each experiment, bacteria were streaked from frozen stocks onto TSB agar plates containing streptomycin (100 µg/mL) and incubated overnight at 37 °C. A fresh plate was prepared for each experiment. Single colonies were used to inoculate 3 mL of TSB supplemented with catalase (100,000 units; Sigma, C40) and grown overnight at 37 °C without shaking. The following morning, cultures were diluted 1:2 in fresh TSB containing catalase and incubated for an additional 4 h, reaching an optical density of approximately OD_600_ = 0.4. Bacterial cultures were then concentrated by centrifugation and resuspended in phosphate-buffered saline (PBS) to a final concentration corresponding to OD_600_ = 2.4. Suspensions were kept on ice prior to loading into injection needles. For infection experiments, approximately 80,000 colony-forming units (CFUs) per individual were injected into the abdominal region using a Hamilton syringe.

### 2.8. In vitro bactericidal activity of hemolymph

Collected hemolymph (15 µl) was diluted to a final volume of 150 µl in Tryptic Soy Broth (TSB), and hemocytes were removed by centrifugation. Cell-free hemolymph was subsequently mixed with *Streptococcus pneumoniae* cultures at a dilution corresponding to a final concentration of approximately 350 CFUs per 10 µl of the final reaction mixture. Hemolymph–bacteria suspensions, together with control reactions lacking hemolymph, were incubated for 2 h at 37 °C. Following incubation, samples were plated onto TSB agar plates containing streptomycin as a selective antibiotic. Bactericidal activity of hemolymph was quantified as a reduction in CFU counts in samples containing hemolymph relative to control samples.

### 2.9. Survival analysis

For survival analysis, experimental individuals were individually injected with a bacterial culture of *Streptococcus pneumoniae* at a dose of 80,000 CFU per individual. This dose was determined to be appropriate for obtaining a survival curve over a period of two to three weeks and was selected based on a preliminary experiment in which different doses were tested.

Injected individuals were housed collectively in breeding containers with a volume of approximately 0.5 L, with about 60 individuals per container. The number of dead individuals was recorded daily, and dead individuals were removed from the containers. The bedding consisting of linden seeds and the water source were regularly replaced at three-day intervals.

### 2.10. Software and statistical analyses

The following software packages were used for data acquisition, analysis, and manuscript preparation. Microscopic images were acquired and initially analyzed using CellSens software (Olympus) and subsequently processed using the open-source software FIJI.

Experimental data were organized and analyzed using GraphPad Prism software, which was also used to perform statistical analyses and generate graphical outputs. Pairwise comparisons between groups were performed using Student’s *t*-test. Comparisons involving multiple groups were analyzed using two-way analysis of variance (two-way ANOVA) followed by Šídák’s multiple-comparisons test. Normality of data distribution was assessed using the Shapiro–Wilk test. Survival data were analyzed using the log-rank (Mantel–Cox) test and the Gehan–Breslow–Wilcoxon test. Final figure assembly and graphical adjustments for publication were performed using Affinity Designer.

## 3. Results

### 3.1. Photoperiod affects both cellular and humoral immunity in P. apterus females

To assess the photoperiod-induces differences in immune system of *P. apterus*, we first analyzed the number of hemocytes present in the hemolymph of males and females maintained at 25°C under reproduction-inducing long-day (LD; 18 h light : 6 h dark) and diapause-inducing SD photoperiodic regimes. Individual cells were visualized by fluorescent labeling of nuclear DNA (DAPI) and actin filaments (phalloidin), and phagocytic activity was assessed using *S. aureus*–pHrodo. This approach enabled reliable discrimination of hemocyte types and quantification of their abundance in hemolymph samples.

Females maintained under SD conditions entered diapause and exhibited a significantly higher number of hemocytes compared with females reared under LD photoperiods, whereas hemocyte numbers in males were comparable across photoperiodic regimes. In addition, females consistently displayed higher hemocyte densities per microliter of hemolymph than males (Fig.1A). Analysis of individual circulating hemocyte populations revealed that these differences were primarily attributable to increased numbers of granulocytes and plasmatocytes (Fig.1B, C). In contrast, the abundance of prohemocytes and the relatively remained unchanged across photoperiodic conditions (Fig.1D). Based on the analysis of phagocytic activity, we conclude that exposure to SD photoperiods enhances the phagocytic capacity of the female immune system even in the absence of a pathogen. Moreover, the increased abundance of granulocytes suggests that humoral immunity may also be proactively upregulated.

**Fig. 1.**
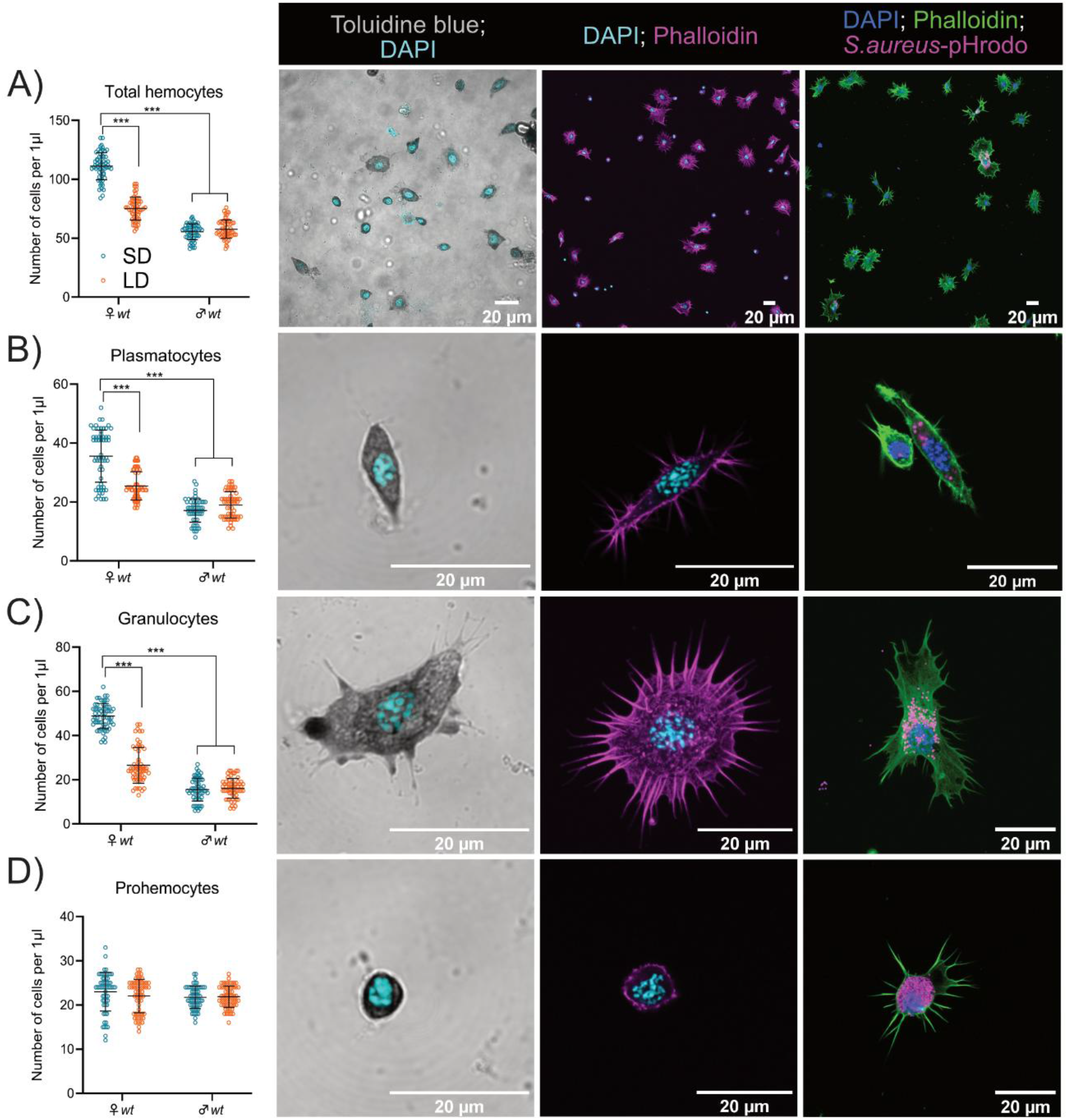
Photoperiod-dependent changes in hemocyte composition in *Pyrrhocoris apterus*. (A) Total hemocyte counts in SD and LD wild-type individuals and representative confocal images of hemocytes stained with Toluidine blue (left), DAP (nucleus) and phalloidin (actin cytoskeleton, magenta; middle), illustrating morphological diversity of hemocyte types. Image on the right illustrates their phagocytic capability following injection of pHrodo™-labeled *Staphylococcus aureus* (magenta). (B)-(D) Quantification of individual hemocyte populations, including plasmatocytes (B), granulocytes (C), and prohemocytes (D) in SD and LD wild-type individuals. Representative confocal images on the left depict the respective hemocyte type stained with Toluidine blue; middle panels show the actin cytoskeleton (phalloidine; magenta), and right panels document phagocytic activity following injection of pHrodo™-labeled *Staphylococcus aureus* particles. Internalized particles appear as magenta puncta. In (A)-(D), ten biological replicates were analyzed, and hemocyte abundance was quantified as the number of cells per µL of hemolymph in six individuals per replicate. Results were compared by two-way ANOVA followed by Šídák’s multiple comparisons test. Values are displayed as mean ± SD; asterisks mark statistically significant differences (***P < 0.001). SD, short-day photoperiod; LD, long day photoperiod.

We therefore extended our analysis to the humoral branch of the immune response. Expression levels of genes encoding antimicrobial peptides (*Defensin, Drosocin*), key components of major immune signaling cascades (*Dorsal-related immunity factor, Relish, Suppressor of cytokine signaling at 36E, spatzle*), and factors involved in the phenoloxidase-mediated melanization response (*Hemiptericin, Peroxidasin*) were quantified. Females maintained under SD conditions exhibited significantly elevated expression of all of these genes (Fig. 2A). Based on insights from *Drosophila* immunology (Neyen et al. 2014), this expression profile suggests a partial activation of the immune system as a whole, rather than activation of a single immune-associated pathway such as Toll or IMD.

**Fig. 2.**
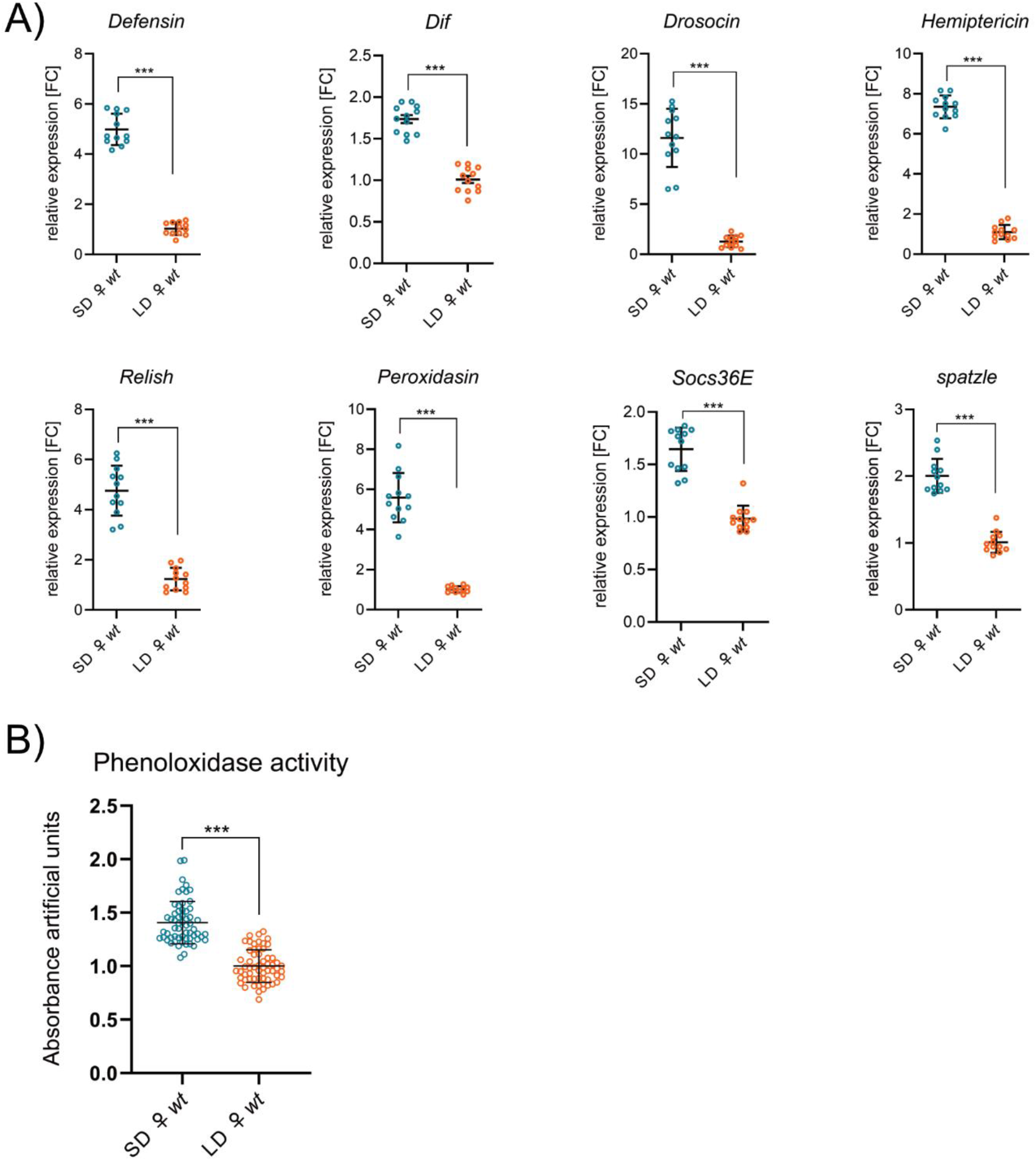
Photoperiod-dependent activation of immunity response in hemocytes of wild-type females. (A) Relative expression of immune-related genes in hemocytes isolated from LD and SD wild-type females. Genes include antimicrobial peptides and immune signaling components. Expression levels were normalized to rp49 and are represented as fold change relative to LD levels arbitrarily set to 1. The experiment was performed in four biological replicates. Each sample consisted of hemolymph pooled from six individuals, and three such samples were analyzed per replicate. (B) Phenoloxidase activity in hemolymph of females reared under LD and SD conditions. The experiment was performed in ten biological replicates, with six individuals analyzed in each replicate. In (A)-(B), results were compared using Student’s t-test and values are displayed as mean ± SD; asterisks mark statistically significant differences (***P < 0.001). *Socs36E, Suppressor of cytokine signaling at 36E*; *Dif, Dorsal-related immunity factor*; SD, short-day photoperiod; LD, long day photoperiod.

Although we also observed an increase in these genes in SD males compared with males maintained under LD conditions, overall, the basal activity of these factors was much higher in females regardless of photoperiod (Supplementary figures, Fig. S1).

Consistent with the transcriptional data, SD females also displayed increased enzymatic activity of prophenoloxidases, closely mirroring the expression patterns of key components of the melanization pathway (Fig. 2B).

### 3.2. Mammalian-type cryptochrome is required for photoperiod-induced changes in the immune system

To test whether the observed immune activation was causally linked to photoperiod perception, we conducted analogous analyses in a recently generated *m-cryptochrome* null mutant (allele *m-cry*^*04*^). As previously demonstrated in *P. apterus*, mutation of *m-cry* prevents entry into diapause even under SD conditions in both sexes (Kotwica-Rolinska et al., 2022, Kaniewska et al., 2024). Comparative analysis of immune parameters in wild-type and *m-cry*^−^*/*^−^ females maintained under SD photoperiods showed that photoperiod-dependent activation of both cellular (Fig. 3A-D) and humoral immunity (Fig. 4A, B) required a functional *m-cry* gene. In males, neither photoperiod nor *m-cry* mutation exerted a pronounced effect on immune parameters (Supplementary Figs. S2 and S3).

**Fig. 3.**
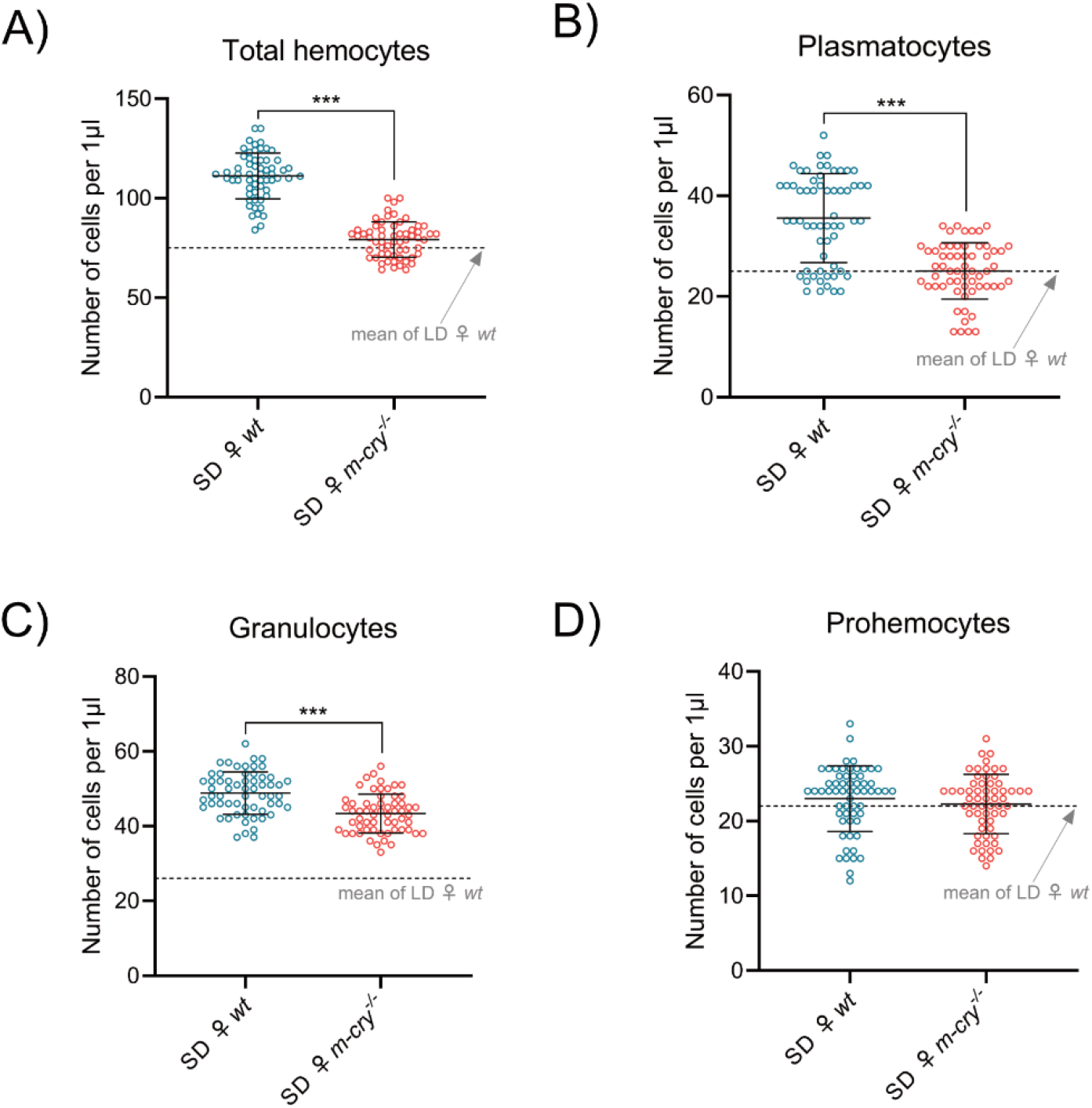
Photoperiod-induced changes in hemocyte numbers require a functional *m-cryptochrome*. (A) Number of total hemocytes in wild-type and *m-cry*^−*/*−^ mutant females reared in SD. (B)-(D) Quantification of individual hemocyte types, including plasmatocytes (B), granulocytes (C), and prohemocytes (D) in wild-type and *m-cry*^−*/*−^ mutant females reared in SD. Ten biological replicates were analyzed, and hemocyte abundance was quantified as the number of cells per µL of hemolymph in six individuals per replicate. In (A)-(D), the experiment was performed in ten biological replicates and the results were compared Student’s t-test. Values are displayed as mean ± SD; asterisks mark statistically significant differences (***P < 0.001). SD, short-day photoperiod; *m-cry, mammalian-type cryptochrome*.

**Fig. 4.**
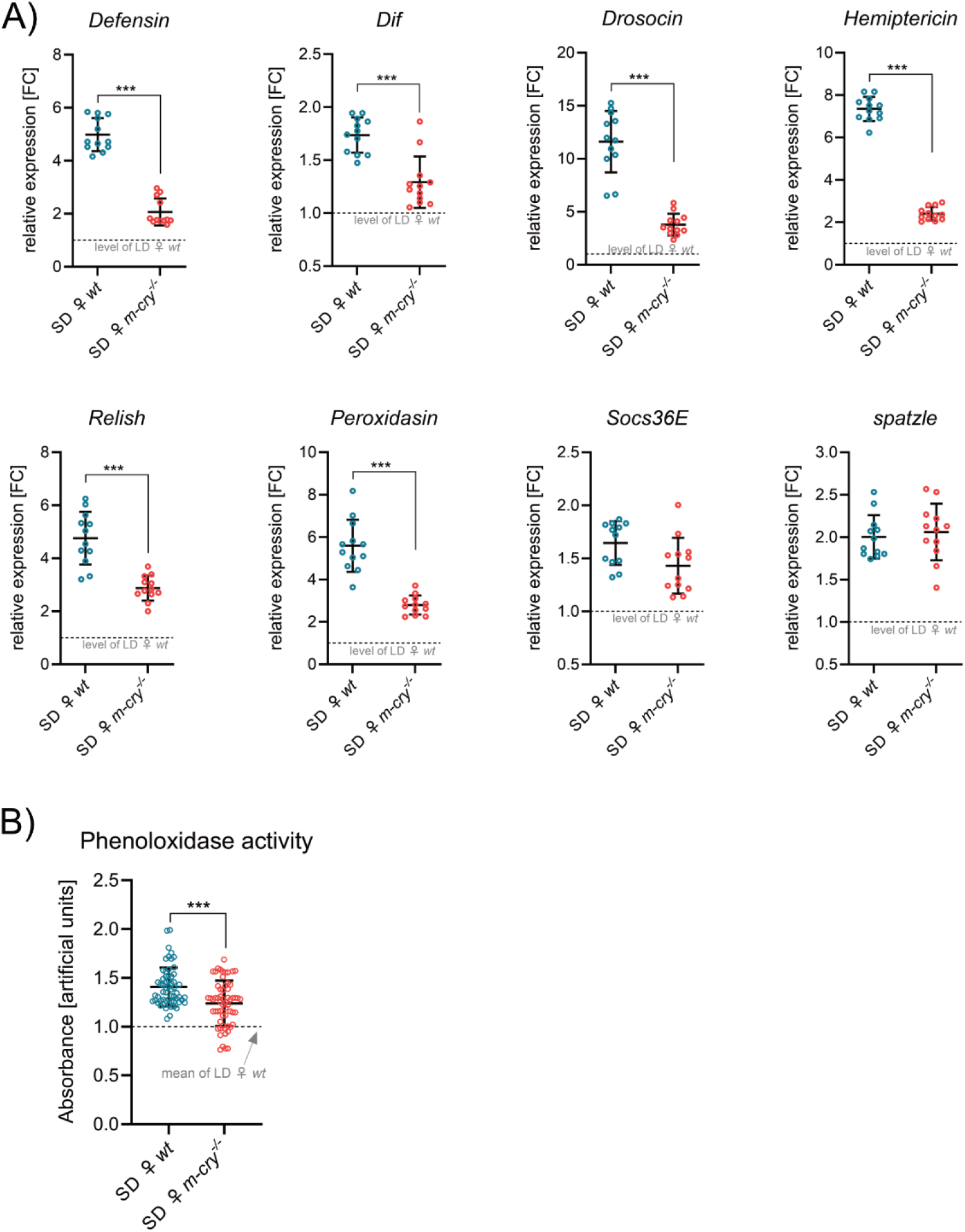
Photoperiod-induced changes in humoral immunity depend on *m-cryptochrome*. (A) Relative expression of immune-related genes in hemocytes isolated from SD wild-type and SD *m-cry*^−*/*−^ mutant females. Expression levels were normalized to rp49 are reported as fold change relative to LD females arbitrarily set to 1. The experiment was performed in four biological replicates. Each sample consisted of hemolymph pooled from six individuals, and three such samples were analyzed per replicate. (B) Phenoloxidase activity in hemolymph of wild-type and *m-cry*^−*/*−^ mutant females reared under SD conditions. The experiment was performed in ten biological replicates, with six individuals analyzed in each replicate. In (A)-(B), results were compared by Student’s t-test and values are displayed as mean ± SD, asterisks mark statistically significant differences (***P < 0.001). SD, short-day photoperiod; *m-cry, mammalian-type cryptochrome*; *Socs36E, suppressor of cytokine signaling at 36E*; *Dif, Dorsal-related immunity factor*.

### 3.3. Photoperiod-induced changes are also observed in natural populations of P. apterus

To assess the ecological relevance of our laboratory findings for the diapause program under complex natural conditions, we collected linden bugs from the field in the vicinity of our research institute at several time points throughout the season. These sampling points corresponded to natural photoperiods comparable to the laboratory LD and SD regimes. Insects collected in June were reproductive, whereas insects collected around the September equinox had entered reproductive diapause. In addition to photoperiod, these linden bugs were exposed to lower autumn temperatures, and all field-collected individuals likely experienced exposure to various naturally occurring pathogens. Field-collected samples showed greater environmental variability and higher inter-individual heterogeneity than laboratory populations. Nevertheless, females collected during periods corresponding to SD conditions consistently exhibited elevated hemocyte numbers, increased expression of immune-related genes, and enhanced prophenoloxidase activity. These patterns closely mirrored those observed under controlled laboratory conditions (Fig. 5A, B, and Supplementary Figs. S3, S4, and S5).

**Fig. 5.**
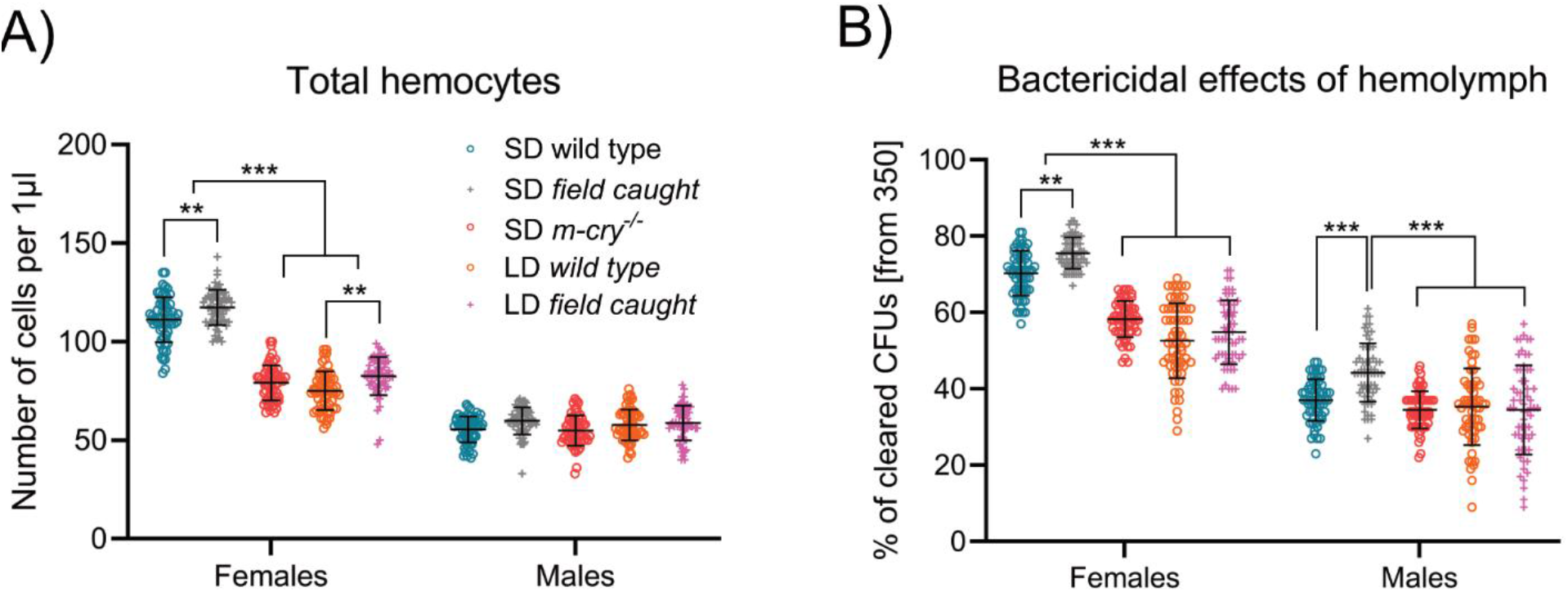
Photoperiod-induced immune priming observed in laboratory animals is also present in field-collected individuals. (A) Total hemocyte counts in SD and LD wild-type individuals, SD-reared *m-cry*^−/−^ mutants and individuals collected in the field at the summer solstice (LD) or the autumnal equinox (SD). Ten biological replicates were analyzed, and hemocyte abundance was quantified as the number of cells per µL of hemolymph in six individuals per replicate. (B) Bactericidal activity of hemolymph from SD and LD wild-type individuals, SD-reared *m-cry*^−*/*−^ mutants, and individuals collected in the field at the summer solstice (LD) or the autumnal equinox (SD). In (B), the experiment was performed in ten biological replicates, with six individuals analyzed in each replicate. Each sample consisted of hemolymph pooled from six individuals, and three such samples were analyzed per replicate. Results were compared by two-way ANOVA followed by Šídák’s multiple comparisons test. Values are displayed as mean ± SD; asterisks mark statistically significant differences (**P < 0.01; ***P < 0.001). SD, short-day photoperiod; LD, long day photoperiod; *m-cry, mammalian-type cryptochrome*.

### 3.4. Photoperiod-induced immune priming enhances resistance to infection

Finally, we asked whether this immune configuration confers a functional advantage during pathogen challenge. Groups of males and females maintained under LD or SD conditions were injected with 80,000 units of *Streptococcus pneumoniae*. Females maintained under LD conditions exhibited a median time to death of 10 days post-infection, whereas females maintained under diapause-inducing SD conditions displayed markedly increased resistance, with a median survival of approximately 17 days (Fig. 6A). These results were consistent across three independent infection experiments. In contrast, males showed only very mild differences in survival between photoperiodic regimes (Fig. 6B), and females were consistently more resistant than males regardless of photoperiod (Fig. 6C, D).

**Fig. 6.**
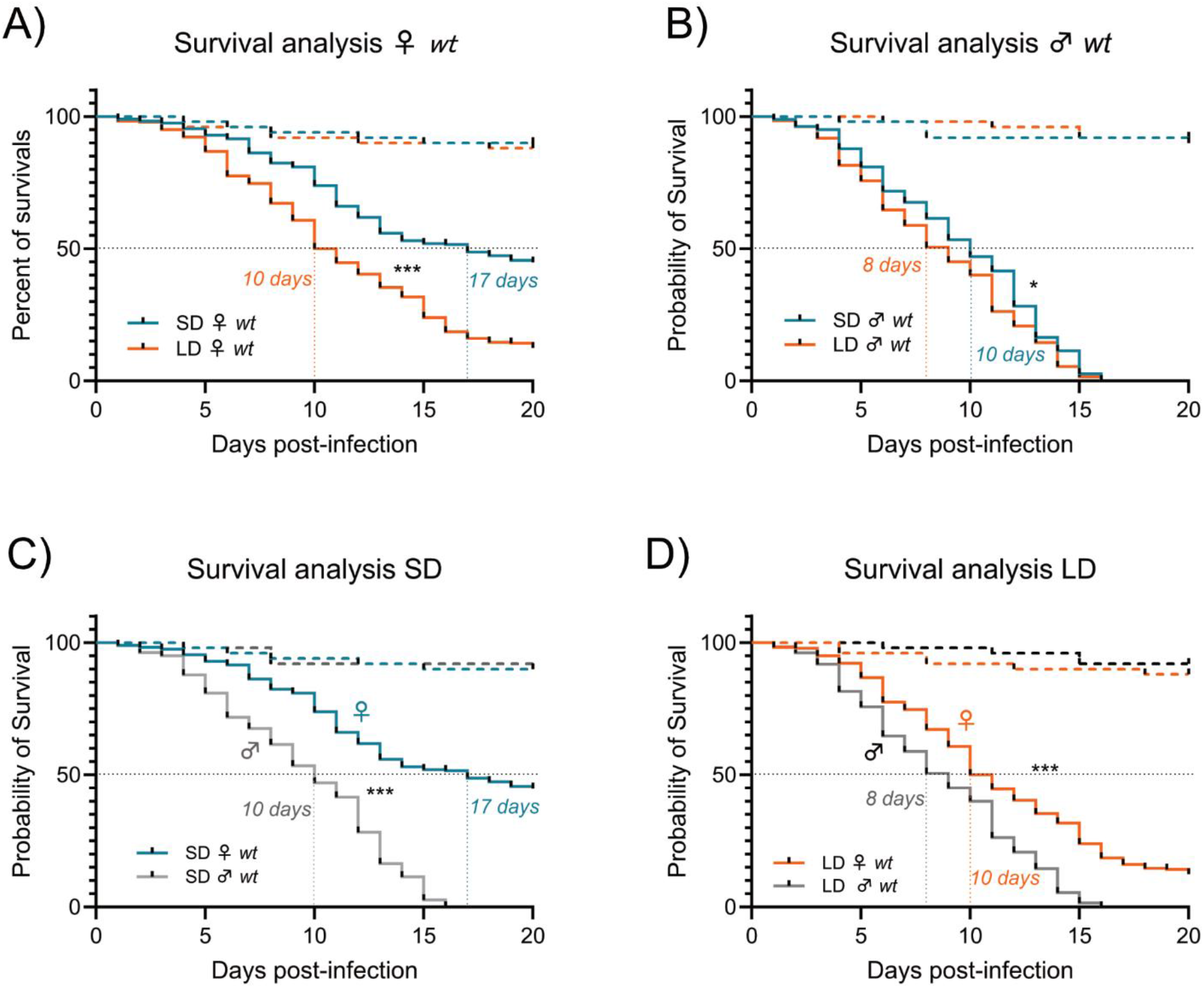
Photoperiod-induced immune priming enhances resistance to bacterial infection. (A)-(D) Kaplan–Meier survival analyses following *Streptococcus pneumoniae* infection in SD- and LD-reared wild-type females (A), SD- and LD-reared wild-type males (B), SD males and SD females (C), and LD males and LD females (D). Results were analyzed by Log-rank and Gehan-Breslow Wilcoxon tests. Dotted lines depict medium time to death. Survival curves of the respective uninfected controls are shown as dashed lines. Three independent experiments were performed and combined into each survival curve, representing at least 150 individuals. Asterisks mark statistically significant differences (*P < 0.05; ***P < 0.001). SD, short-day photoperiod; LD, long day photoperiod.

These differences in resistance to infection were supported by *in vitro* assays assessing the bactericidal activity of cell-free hemolymph. Incubation of *S. pneumoniae* with isolated hemolymph significantly reduced bacterial viability, with the strongest effect observed in hemolymph derived from SD females (Fig. 5B).

Collectively, our results demonstrate that photoperiod has a profound and sex-specific impact on immune system organization and function in *P. apterus*. SD, diapause-inducing conditions are associated with coordinated activation of both cellular and humoral immunity in females. This response is manifested by increased total hemocyte numbers, driven primarily by elevated plasmatocyte and granulocyte populations, upregulation of immune-related genes, enhanced prophenoloxidase activity, and improved resistance to bacterial infection. These immune changes are functionally linked to photoperiod-dependent programs, as they require a functional *m-cry* gene. Comparable female-specific immune activation was also observed in field-collected individuals under natural SD conditions, despite greater environmental variability. In contrast, males exhibited largely photoperiod-independent immune parameters across all assays. Together, these findings reveal pronounced intersexual differences in immune responsiveness to photoperiod and identify day-length perception as a key regulator of immune preparedness in females entering diapause.

## 4. Discussion

Our data show that photoperiod is an important regulator of immune system in *Pyrrhocoris apterus*. SD conditions, which induce reproductive diapause, were associated with anticipatory activation of both cellular and humoral immunity in females. This effect was mechanistically linked to photoperiodic signaling, because in the absence of functional *m-cryptochrome*, a key component required for proper photoperiodic responses, immune changes in SD females were largely lost. Importantly, the same overall pattern was also observed in field-collected individuals exposed to naturally occurring LD and SD conditions, where the effects of seasonal state on measured immune parameters closely resembled those observed under laboratory conditions.

### 4.1. Short-day females exhibit broad immune preparedness

Our results show that females maintained under SD conditions have increased numbers of granulocytes and plasmatocytes together with elevated expression of humoral immune factors. This pattern is consistent with the observation that these females display enhanced resistance to experimental bacterial infection and that their hemolymph exerts stronger bactericidal activity in vitro. Taken together, these findings indicate that SD females enter a broadly enhanced state of immune preparedness rather than showing an isolated change in a single immune trait.

At first sight, these observations may appear somewhat unexpected in light of previous literature, where diapause has often been associated with reduced cellular immunity (Raina and Bell 1974, Nakamura et al. 2011). However, available studies also indicate that seasonal immune regulation is highly species-specific (Ferguson and Sinclair 2017). In this context, it may be particularly relevant that *P. apterus* is a hemimetabolous insect. Much of the existing literature on diapause and immunity concerns larval stages of holometabolous species, in which hemocyte abundance and differentiation are additionally shaped by hematopoiesis, tissue remodeling, and metamorphosis (Hultmark and Ando 2022, Krejcova et al. 2024). Seasonal immune regulation in adult hemimetabolous insects may therefore follow a substantially different logic.

Our data further point to an intriguing relationship between reproductive state and immunity in females. In *m-cryptochrome* mutants, whose disruption abolishes diapause induction under SD conditions (Kaniewska et al., 2024; Kotwica-Rolinska et al. 2022), both cellular and humoral immune activity remained at levels comparable to those of long-day wt females. This pattern raises the possibility that reproductive activation and immune preparedness are negatively coupled, potentially through competition for shared systemic resources or through common endocrine regulation. At the same time, immune cells are not merely defensive effectors, but major immunometabolic regulators capable of influencing systemic metabolism and reproductive physiology (Bajgar et al., 2021, Krejcova et al. 2023). Future work will therefore need to clarify whether the observed inverse relationship between ovarian maturation and immune activation reflects direct resource competition, hormonal coordination, or bidirectional communication between reproductive and immune tissues.

### 4.2. Seasonal immune preparedness is strongly sex-specific in P. apterus

A particularly striking finding of this study is that photoperiod-dependent immune changes were observed almost exclusively in females. Comparable sex-specific seasonal immune effects have not yet been clearly established in other insect systems, but sex-specific strategies for surviving unfavorable seasons are by no means unusual (Schebeck et al., 2024, Meuti et al. 2024). In *P. apterus*, this observation is especially interesting in light of the work of Hejníková and colleagues (Hejníkova et al. 2022), who showed that environmental cues controlling diapause entry and termination differ substantially between males and females. These findings suggest that the downstream physiological programs triggered by photoperiod are also sexually dimorphic, and our data indicate that immunity forms part of this dimorphism. It will be important in future work to determine whether male immune system responds only to a broader or more complex set of seasonal cues than day length alone. Similarly, it will be interesting to characterize immune response during P. apterus aestivation, which seem to be connected to insulin signaling (Smykal et al., 2020).

### 4.3. Functional m-cryptochrome is required for photoperiodic immune regulation

Our experiments clearly show that mutation of *m-cryptochrome*, which disrupts proper photoperiodic signaling and prevents reliable diapause induction, also abolishes the seasonal immune phenotype seen in wild-type females. This observation is important because it links immune regulation directly to the photoperiodic response pathway rather than to a nonspecific correlation of seasonal state. In this sense, the immune phenotype appears to be integrated into the same central seasonal program that governs diapause induction (Kotwica-Rolinska et al. 2017, Kotwica-Rolinska et al. 2022).

Previous work in *P. apterus* has shown that the photoperiodic timer is linked to endocrine output from corpora allata, with juvenile hormone acting as a major coordinator of diapause-related physiological change (Hejnikova et al., 2016, 2022; Bajgar et al., 2013a; Urbanova et al., 2016). It is therefore plausible that hormonal signals downstream of central photoperiodic time measurement also participate in regulating immune function. However, this remains to be tested directly. The *P. apterus* system offers attractive opportunities for such experiments, including manipulation of juvenile hormone signaling through hormone analogs or targeted disruption of key signaling components.

At the same time, we cannot exclude the contribution of gene pleiotropy. In *m-cryptochrome* mutants, not only central photoperiodic signaling but also clock-related processes in peripheral oscillators are likely to be affected (Bajgar et al. 2013, Smykal et al. 2014). Similarly, noncanonical action of circadian clock genes was reported for diapause entry in beetle *Harmonia axyridis* (Gao et al., 2025). The observed immune phenotype may therefore reflect a combination of disrupted photoperiodic measurement and altered organization downstream of this device. Dissecting these contributions will require further targeted experiments.

### 4.4. Ecological relevance of seasonal immune preparedness

One major strength of the *P. apterus* model is that controlled laboratory experiments can be combined with repeated sampling of natural populations in the vicinity of the research institute. To test whether the laboratory phenotype has ecological relevance, we collected field individuals under natural photoperiodic conditions approximating the long-day and SD regimes used in the laboratory. Despite the much greater environmental variability experienced by wild populations, the main differences in immune system organization in SD females remained clearly detectable. This strongly supports the view that the observed immune phenotype is not a laboratory artifact but represents a biologically meaningful seasonal adaptation.

Future work should examine immune activity across the full annual cycle, including winter and early spring populations, where seasonal history, cold exposure, and fluctuating environmental conditions may further shape immune state. It will also be valuable to integrate immune measurements with analyses of microbiome composition, natural pathogen burden, and other ecological variables. Such studies will help place seasonal immune regulation in *P. apterus* into its full ecological context.

## 5. Conclusion

Our results identify photoperiod as a central organizer of immune system function in *P. apterus*. SD, diapause-inducing conditions trigger a robust, female-specific immune preparedness program that coordinately enhances cellular and humoral immunity, leading to increased infection resistance. This response strictly depends on functional Cryptochrome-mediated day-length perception and is absent in photoperception-deficient mutants, underscoring a direct link between environmental sensing and immune regulation. The persistence of this pattern in field-collected females highlights its ecological relevance and reveals pronounced intersexual divergence in how immunity is integrated into seasonal life-history strategies.

## 6. Acknowledgement

The authors would like to thank Hanka Vaněčková and Lucie Hrádková for their excellent technical assistance and dedicated care of experimental insect cultures, as well as for their essential support in the daily operation of the laboratory. The authors used services of the Czech-BioImaging research infrastructure, specifically the Laboratory of Microscopy and Histology, BC CAS, supported by project LM2023050 funded by the Ministry of Education, Youth and Sports of the Czech Republic, with the instrumental equipment co-financed by the European Union. GK, AB and microscopy work were supported by the project reg. no. CZ.02.01.01/00/23_021/0009529 (iBEE – Intelligent Beekeeping: Modern Biotechnologies for Healthy Bees), co-funded by the European Union. VS, DD, and the transcriptomic part of the study were supported by the Ministry of Education, Youth and Sports, project LUAUS25273.

## CRediT authorship contribution statement

Categories: Conceptualization – DD, AB; Data curation – GK, VS, AB; Formal analysis – GK, VS, DD, AB; Funding acquisition – DD, AB; Investigation – GK, VS, DD, AB; Methodology – GK, VS, AB; Project administration – DD, AB; Resources – GK, VS, DD, AB; Supervision – DD, AB; Validation – GK, VS, DD, AB; Visualization – GK, VS, DD, AB; Writing – original draft – GK, DD, AB; Writing – review and editing – GK, VS, DD, AB

