## Supplementary material: for "Photoperiod induces sex-specific immune priming in *Pyrrhocoris apterus*"

Short title: Immune priming in diapause

Authors: Adam Bajgar[a,b,1,*], Gabriela Krejčová[a,b,1], Vlastimil Smýkal[b], and David Doležel[b,**]

[a] *Department of Molecular Biology and Genetics, Faculty of Science, University of South Bohemia, 370 05 České Budějovice, Czech Republic.*
[b] *Institute of Entomology, Biology Centre of the Czech Academy of Sciences, 370 05 České Budějovice, Czech Republic.*

[1]These authors contributed equally to this work
* Corresponding author
** Corresponding author

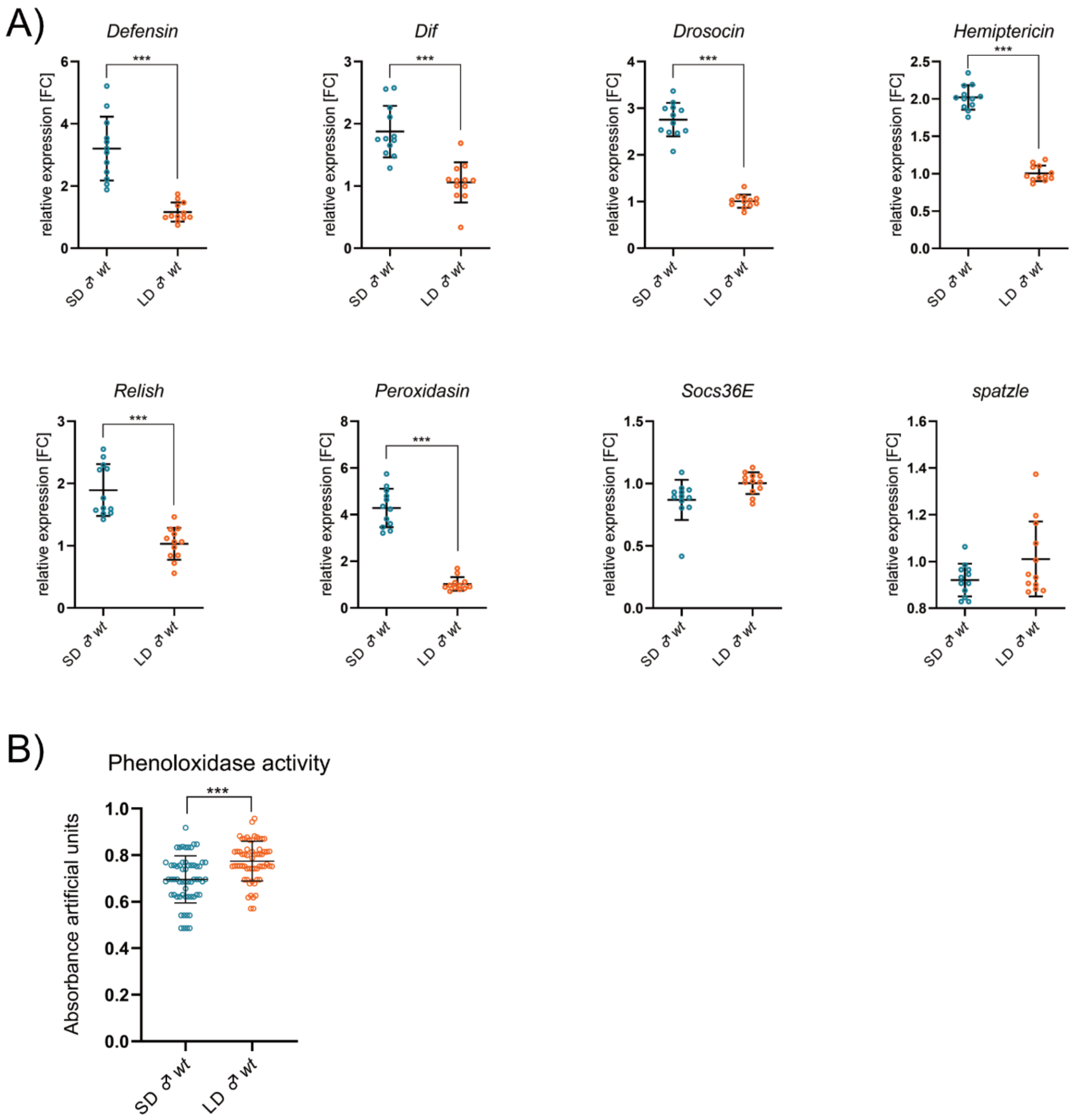


**Supplementary Fig. 1.** (A) Relative expression of immune-related genes in hemocytes isolated from LD and SD wild-type males. Genes include antimicrobial peptides and immune signaling components. Expression levels were normalized to rp49 and are represented as fold change relative to LD levels arbitrarily set to 1. The experiment was performed in four biological replicates. (B) Phenoloxidase activity in hemolymph of males reared under LD and SD conditions. The experiment was performed in ten biological replicates. In (A)-(B), results were compared using Student's t-test and values are displayed as mean ± SD; asterisks mark statistically significant differences (***$P < 0.001$). *Socs36E*, *Suppressor of cytokine signalling at 36E*; *Dif*, *Dorsal-related immunity factor*; SD, short day photoperiod; LD, long day photoperiod.

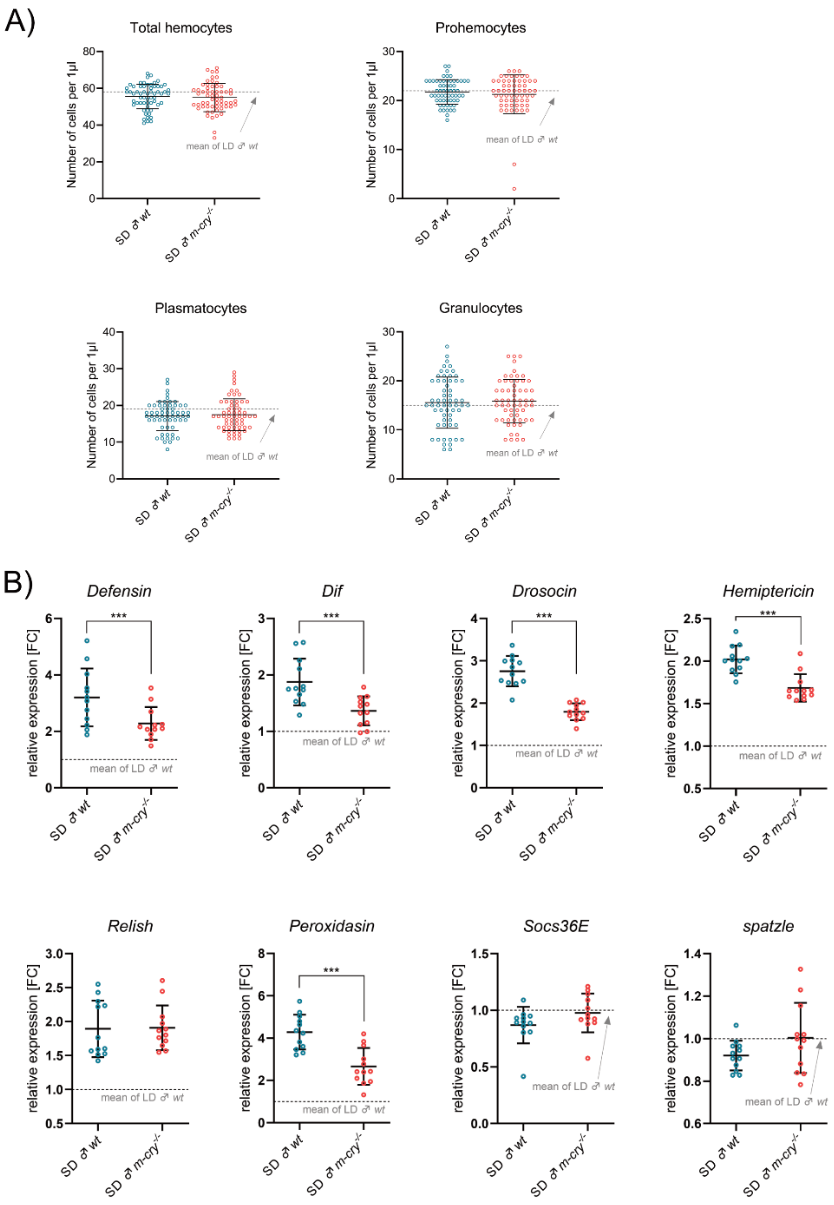


**Supplementary Fig. 2.** (A) Number of total hemocytes and individual hemocyte types, including prohemocytes plasmatocytes, and granulocytes in wild-type and *m-cry*$^{-/-}$ mutant males reared in SD. Ten biological replicates were analyzed and the number of hemocytes was quantified as number of cells per μL of hemolymph. (B) Relative expression of immune-related genes in hemocytes isolated from SD wild-type and SD *m-cry*$^{-/-}$ mutant males. Expression levels were normalized to rp49 are reported as fold change relative to LD wild-type males arbitrarily set to 1. The experiment was performed in four biological replicates. In (A)-(B), results were compared by Student's t-test. Values are displayed as mean ± SD; asterisks mark statistically significant differences (***P < 0.001). SD, short day photoperiod; *m-cry*, *mammalian-type cryptochrome*; *Socs36E*, *Suppressor of cytokine signalling at 36E*; *Dif*, *Dorsal-related immunity factor*.

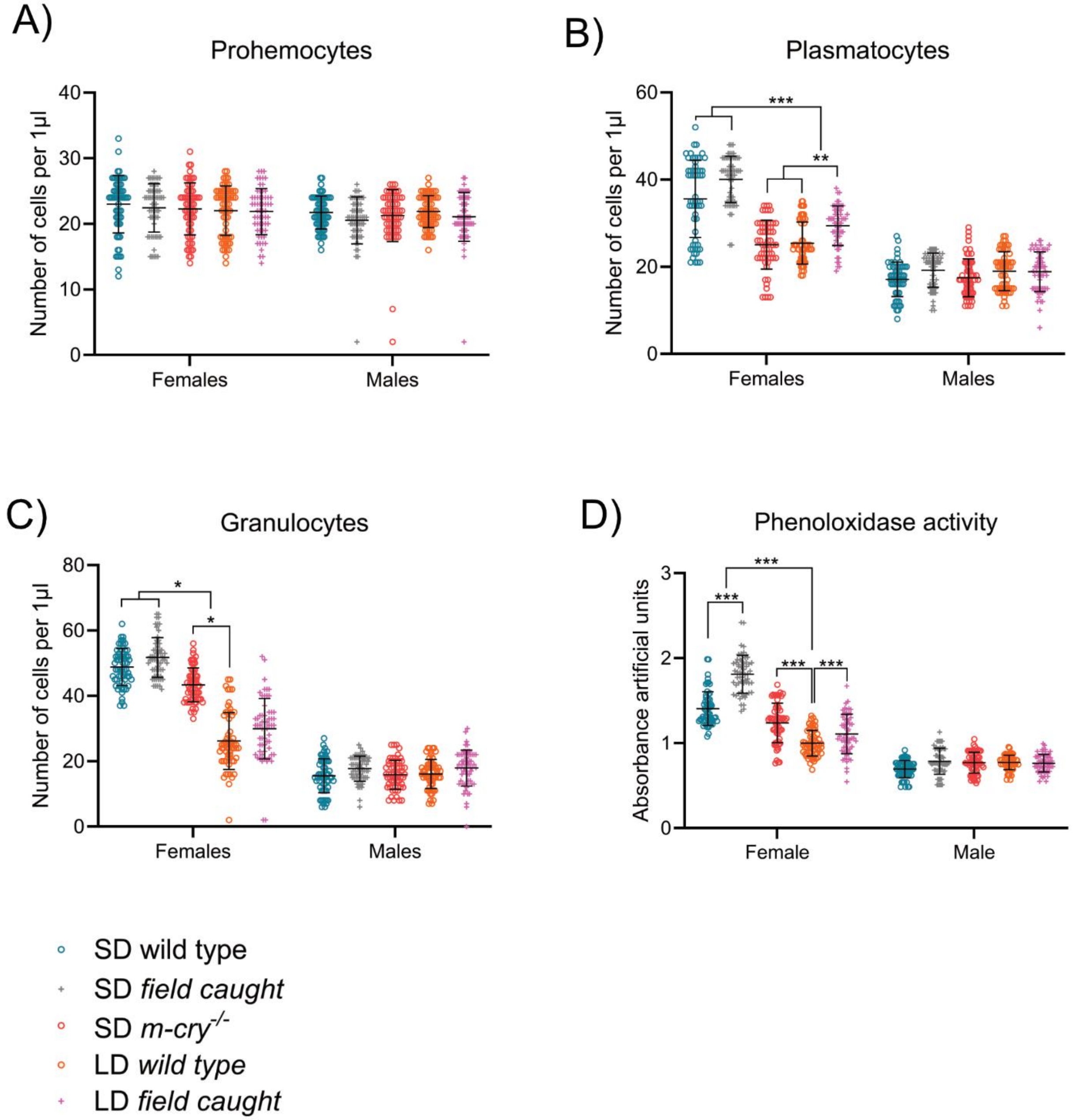


**Supplementary Fig. 3.** (A)-(C) Quantification of individual hemocyte populations, including prohemocytes (A), plasmatocytes (B), and granulocytes (C) in SD and LD wild-type individuals, SD-reared *m-cry*$^{-/-}$ mutants and individuals collected in the field at the summer solstice (LD) or the autumnal equinox (SD). Ten biological replicates were analyzed and the number of hemocytes was quantified as number of cells per µL of hemolymph. (D) Phenoloxidase activity in hemolymph of SD and LD wild-type individuals, SD-reared *m-cry*$^{-/-}$ mutants and individuals collected in the field at the summer solstice (LD) or the autumnal equinox (SD). The experiment was performed in ten biological replicates. In (A)-(D), results were compared by two-way ANOVA followed by Šídák's multiple comparisons test. Values are displayed as mean ± SD, asterisks mark statistically significant differences (*P < 0.05; **P < 0.01; ***P < 0.001). SD, short day photoperiod; *m-cry*, *mammalian-type cryptochrome*.

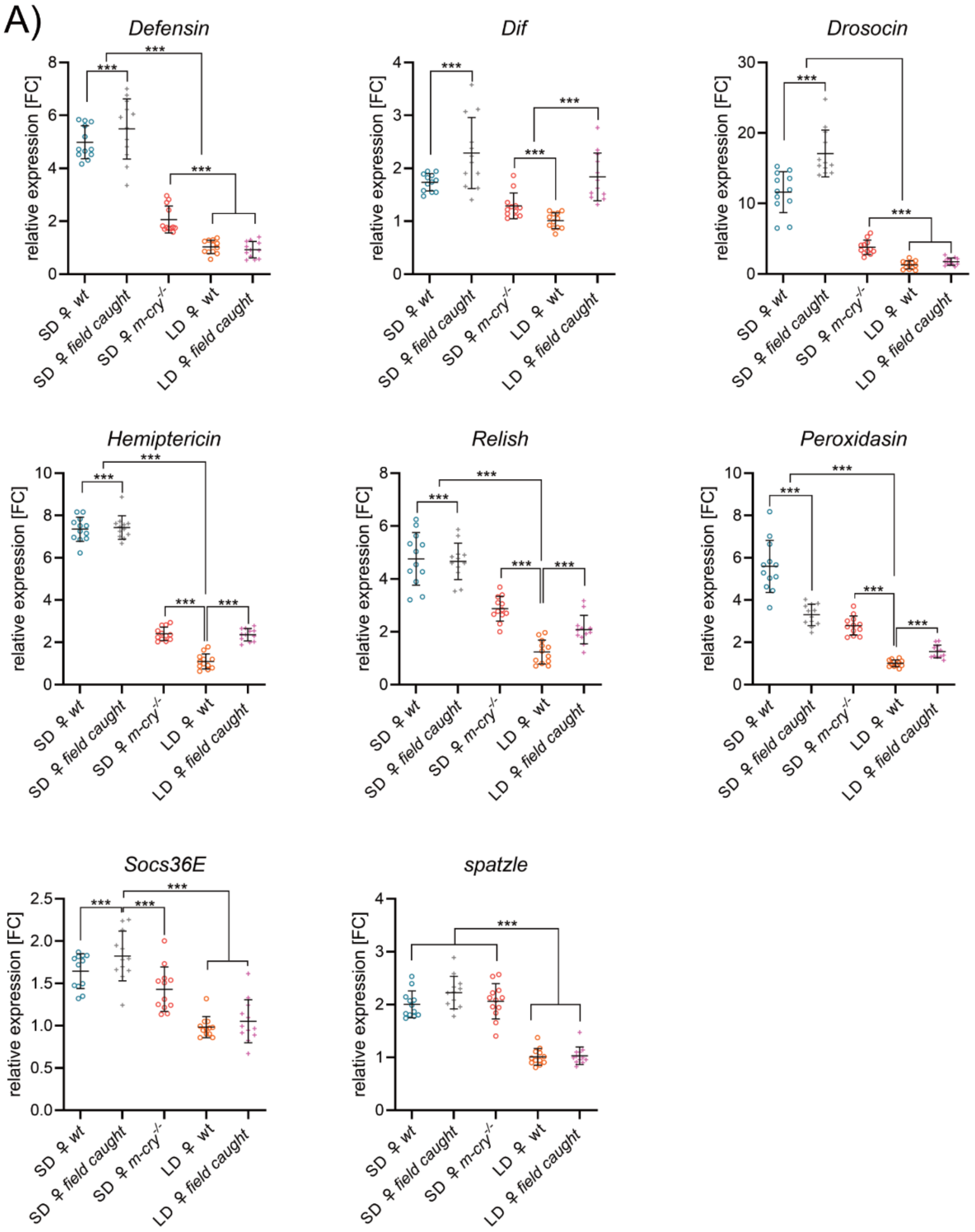


**Supplementary Fig. 4.** (A) Relative expression of immune-related genes in hemocytes isolated from SD and LD wild-type females, SD-reared *m-cry*$^{-/-}$ mutant females and females collected in the field at the summer solstice (LD) or the autumnal equinox (SD). Expression levels were normalized to rp49 are reported as fold change relative to LD females arbitrarily set to 1. The experiment was performed in four biological replicates. Results were compared by two-way ANOVA followed by Šídák's multiple comparisons test. Values are displayed as mean ± SD, asterisks mark statistically significant differences (***$P < 0.001$). SD, short day photoperiod; *m-cry*, *mammalian-type cryptochrome*; *Socs36E*, *Suppressor of cytokine signalling at 36E*; *Dif*, *Dorsal-related immunity factor*.

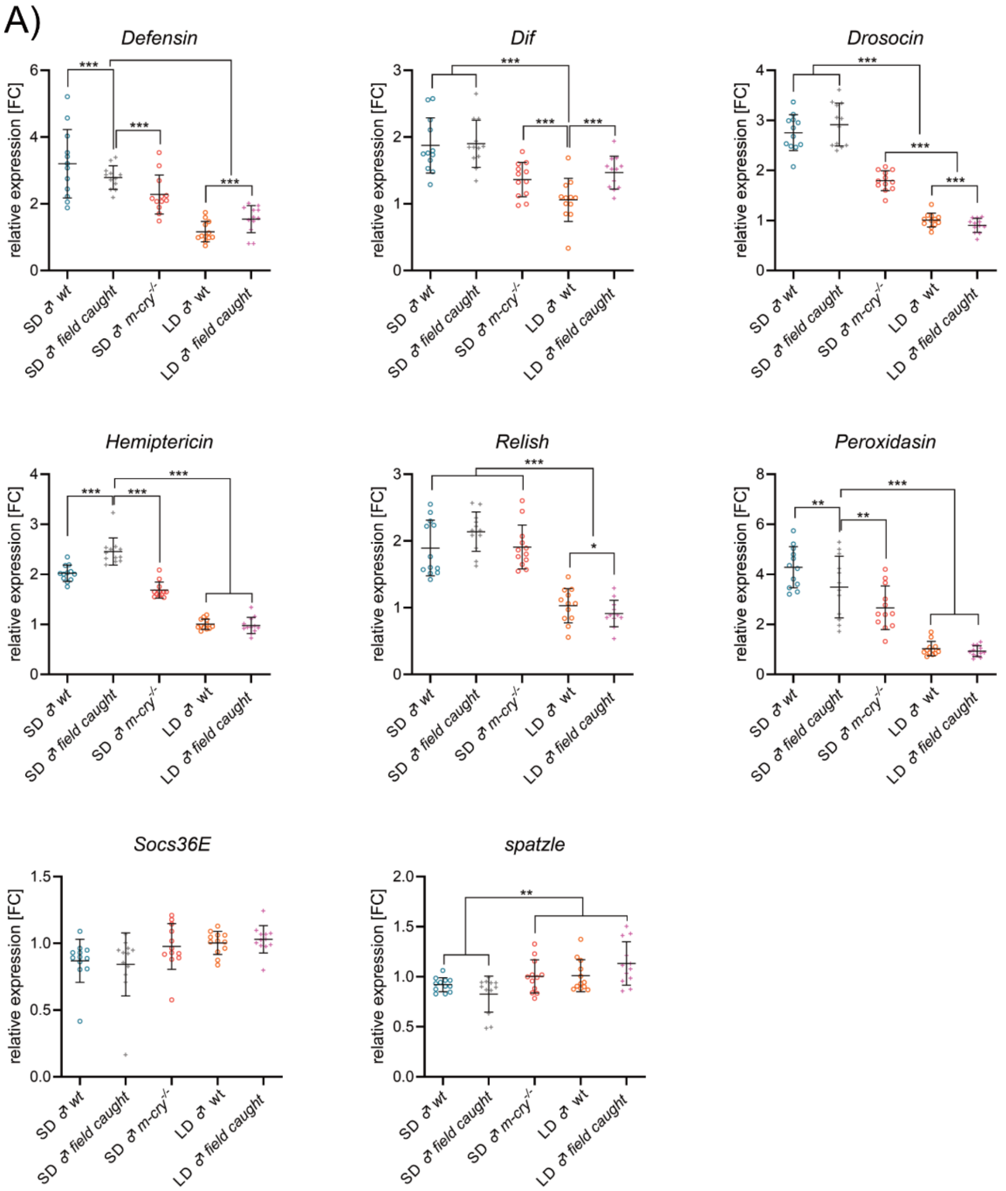


**Supplementary Fig. 5.** (A) Relative expression of immune-related genes in hemocytes isolated from SD and LD wild-type males, SD-reared *m-cry*$^{-/-}$ mutant males and males collected in the field at the summer solstice (LD) or the autumnal equinox (SD). Expression levels were normalized to rp49 are reported as fold change relative to LD females arbitrarily set to 1. The experiment was performed in four biological replicates. Results were compared by two-way ANOVA followed by Šídák's multiple comparisons test. Values are displayed as mean ± SD, asterisks mark statistically significant differences (**P < 0.01; ***P < 0.001). SD, short day photoperiod; *m-cry*, *mammalian-type cryptochrome*; *Socs36E*, *Suppressor of cytokine signalling at 36E*; *Dif*, *Dorsal-related immunity factor*.

A

B

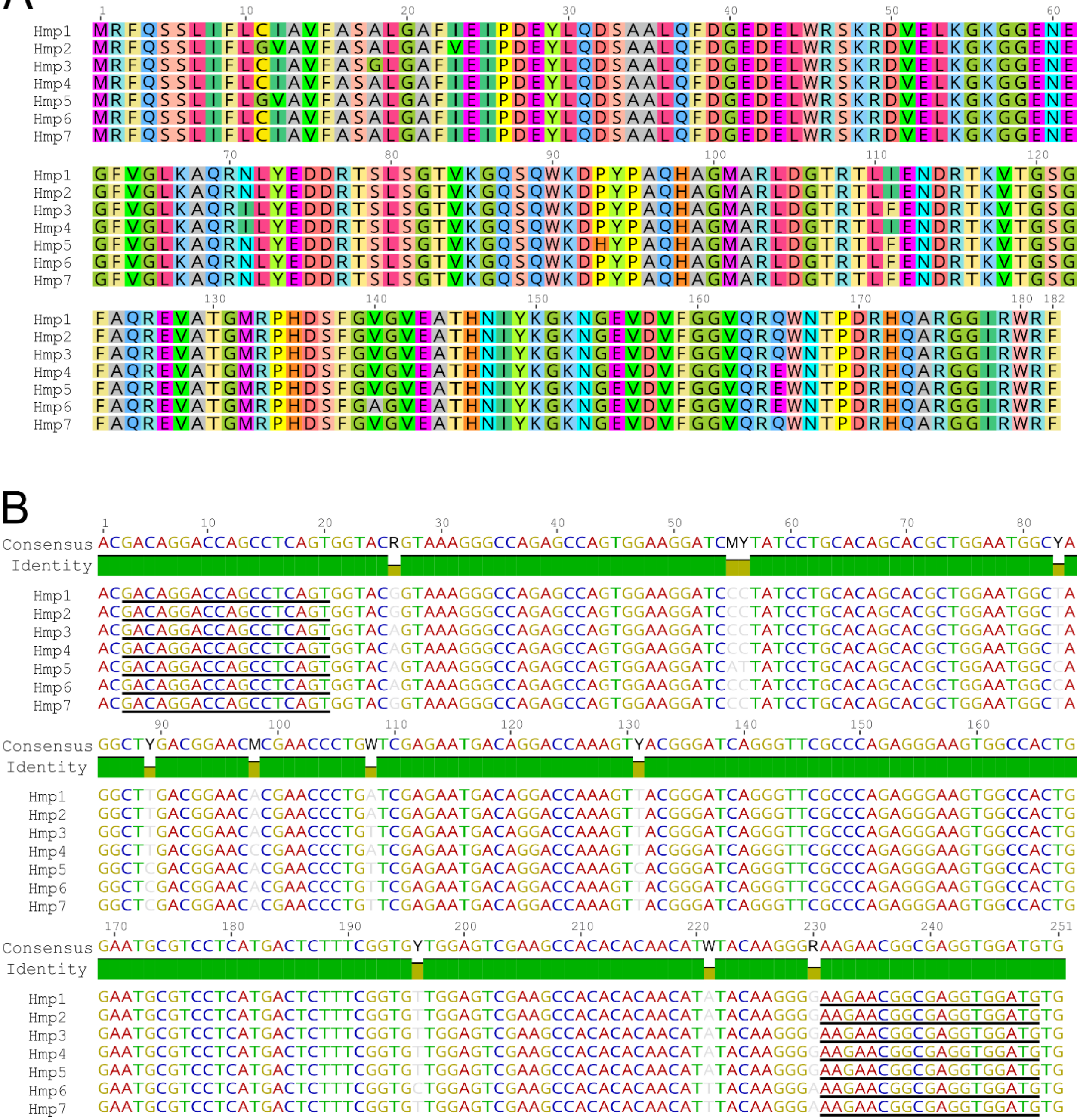

**Supplementary Fig. 6.** (A) Protein and mRNA alignments of *P. apterus* hemiptericins. The underlined sequences in panel B refer to the forward and reverse primers, respectively. The IUPAC nucleotide code is used to represent degenerate positions in amplicons.

```
LOCUS       Defensin                 526 bp    DNA     linear   UNA 03-MAR-2026
DEFINITION  Pyrrhocoris apterus Defensin, complete mRNA.
ACCESSION
KEYWORDS    .
SOURCE      Pyrrhocoris apterus
  ORGANISM  Pyrrhocoris apterus
            cellular organisms; Eukaryota; Opisthokonta; Metazoa; Eumetazoa;
            Bilateria; Protostomia; Ecdysozoa; Panarthropoda; Arthropoda;
            Mandibulata; Pancrustacea; Hexapoda; Insecta; Dicondylia;
            Pterygota; Neoptera; Paraneoptera; Hemiptera; Prosorrhyncha;
            Heteroptera; Euheteroptera; Neoheteroptera; Panheteroptera;
            Pentatomomorpha; Pyrrhocoroidea; Pyrrhocoridae; Pyrrhocoris.
FEATURES             Location/Qualifiers
     5'UTR           1..58
                     /gene="Def"
     gene            1..526
                     /gene="Def"
     mRNA            join(1..143,144..170,171..526)
                     /gene="Def"
     source          1..>526
                     /mol_type="mRNA"
                     /organism="Pyrrhocoris apterus"
                     /db_xref="taxon:37000"
     CDS             join(59..143,144..170,171..325)
                     /gene="Def"
                     /product="Defensin"
                     /translation="MKFVVLFIFTVVVAMASAHPYIPVDEDADVPDAIPEEYHSLRVK
                     RATCDVLSFSSKWFTPNHSACAIHCIAKGYKGGSCKKAICHCRR"
                     /codon_start=1
     primer_bind     138..157
                     /standard_name="Def-Fwd"
     primer_bind     complement(273..291)
                     /standard_name="Def-Rev"
     3'UTR           326..526
                     /gene="Def"
ORIGIN
        1 agctagagtc cagcagtcta caggtatcac agccaattct aacaagaaac caacaacaat
       61 gaaatttgta gttctcttta tcttcacagt agttgtagca atggcctcag cacatcccta
      121 cattccggtt gacgaggatg ctgatgttcc tgatgccata ccagaagaat atcacagcct
      181 ccgtgtgaag agggcaacat gcgacgtatt gagtttttcc tccaagtggt tcactccaaa
      241 ccactccgca tgcgccatcc attgcatcgc caaaggctac aagggaggaa gctgcaaaaa
      301 agctatctgc cactgcagga gataaacagc aaaaacccaa aatacaactc aaatacaact
      361 gtatttatac aactcattaa accaatccaa attatctaaa tacaaaagta ttgataaagc
      421 aatattatta atatgttatt aaatgtcagt attataatat atatatattt tacattaaaa
      481 tattatatat cttatattaa gaaataaatc tcattaatac tatata
//
```

```
LOCUS       Dorsal-related immunity factor, isoform A       2017 bp    DNA     linear
UNA 03-MAR-2026
DEFINITION  Pyrrhocoris apterus Dorsal-related factor, isoform A, complete
            mRNA.
ACCESSION
KEYWORDS    .
SOURCE      Pyrrhocoris apterus
  ORGANISM  Pyrrhocoris apterus
            cellular organisms; Eukaryota; Opisthokonta; Metazoa; Eumetazoa;
            Bilateria; Protostomia; Ecdysozoa; Panarthropoda; Arthropoda;
            Mandibulata; Pancrustacea; Hexapoda; Insecta; Dicondylia;
            Pterygota; Neoptera; Paraneoptera; Hemiptera; Prosorrhyncha;
            Heteroptera; Euheteroptera; Neoheteroptera; Panheteroptera;
            Pentatomomorpha; Pyrrhocoroidea; Pyrrhocoridae; Pyrrhocoris.
FEATURES             Location/Qualifiers
     5'UTR           1..335
                     /gene="Dif"
     gene            1..2017
                     /gene="Dif"
     mRNA            join(1..312,313..384,385..505,506..593,594..837,838..981,
                     982..1080,1081..1214,1215..1311,1312..1383,1384..1422,
                     1423..1691,1692..2017)
                     /gene="Dif"
                     /note="Dorsal-related immunity factor, isoform A"
     source          1..2017
                     /mol_type="mRNA"
                     /organism="Pyrrhocoris apterus"
                     /db_xref="taxon:37000"
     CDS             join(336..384,385..505,506..593,594..837,838..981,
                     982..1080,1081..1214,1215..1311,1312..1383,1384..1422,
                     1423..1691,1692..1862)
                     /gene="Dif"
                     /product="Dorsal-related immunity factor, isoform A"
                     /translation="MDFQSDDSNQINISDVIEVIETDEAFGFPQPRRYSSDRGMAGAR
                     LKIIEQPAPKGLRFRYECEGRSAGSLPGISSTPENKTYPTVQIVGYTGKAVIIVSCVT
                     KDPPYKPHPHNLVGRETCKNGVCRVKLLGDTMTATFANLGIQCVKKKDIEEALKVREK
                     LRMDPFGTGFSHKNQTSSIDLNSVRLCFQAFLEGNEKERFTVPLEPVVSEPIYDKKAI
                     ADLIICKISHGAASVAGGQEIIILCEKVAKEDISIRFFEEGRDGVTWEGYADFIPGDV
                     HKQVAISLRTPRYRTLEIEQPVKVYIQLRRPSDGKTSSPHPFLFTPLDSDGSKKRKRQ
                     KYSPNDLILQNLDHMRPETSAMSIKREPLSSPPNNRIVVSPGKVMSPLQARTPSPQYQ
                     QSFLVQGASGVSKTYVSMGQSSGGQLLVPFMLDQQQQQQQTAITNANEIGRFLFNMD
                     TQQTSVEVNQYEVIDHSLSNNLSASLNLIEQDYQPDASSESLTRLTTNAYNQLTNNFP
                     "
                     /codon_start=1
     primer_bind     1105..1122
                     /standard_name="Dif-Fwd"
     primer_bind     complement(1276..1293)
                     /standard_name="Dif-Rev"
     3'UTR           1863..2017
                     /gene="Dif"
ORIGIN
        1 agtctatttc agtactcccg tccgattgct gcactttcca aaaataatct tttcagtttt
       61 cttcttcatg tgtatgattc gactttaagt cttccctttt aacgagaact tttaaaagtg
      121 tccacaatat attatttaat atatttaata atcgagtgtt actgtgttta tatattcttt
      181 aaagtatata tataatcagt tgataataac tgttgtcaac ttatttattt ctgagaagtc
      241 cttgagtatt tatataaata ctcttttta ttaaaacaat ttcaattcta acagaatgtg
      301 gaaagtatag agatactata cgttatgaaa gaaacatgga tttccaaagt gacgacagta
      361 accaaattaa tataagtgat gtcattgaag tcatcgagac agatgaagct ttcggctttc
      421 cccagccaag aagatatagt tcggacagag gaatggctgg cgcacgactt aaaattattg
      481 aacagcctgc tccgaaagga ctaaggttta gatatgaatg cgaaggtaga tccgctggca
      541 gtttgccggg tatcagttcg actccagaaa ataaaacata tccaacagtg cagattgttg
      601 gatacaccgg caaagcagta ataatcgtct cgtgcgttac aaaggatccg ccttataagc
      661 cgcatcctca caacctcgtc ggtagagaga cgtgtaaaaa cggagtatgt agggtgaaat
```

```
  721 tactcggtga cacgatgact gcaacattcg ccaatttagg aatacagtgc gtaaagaaga
  781 aagatataga agaggcgcta aaggtcagag aaaaactaag gatggatccg tttgggactg
  841 gattcagtca taaaaatcaa acgagttcaa ttgacctcaa ttctgtaaga ttatgctttc
  901 aagcattcct cgaaggtaat gagaaggaga gattcaccgt tcctttagag ccagtggttt
  961 cagaaccgat atacgataaa aaagccatcg ctgatctcat tatttgcaag atcagtcatg
 1021 gggcagcatc agttgcaggc ggccaggaaa ttattatcct ttgtgagaaa gtagctaaag
 1081 aagacattag catacgattt ttcgaggaag gacgagacgg agttacttgg gaaggttatg
 1141 cggatttcat tcccggcgat gtacataaac aagtggctat ctctttacgc acccctcgct
 1201 atcgaaccct tgagattgag caaccggtca aagtgtatat tcagctgaga agaccttcgg
 1261 acggtaaaac cagttctccg caccctttcc tgtttactcc actggattca gatgggtcca
 1321 aaaagaggaa aaggcaaaaa tatagcccta acgatttgat attacaaaac ctggatcata
 1381 tgagacctga aactagtgct atgtccatca aacgagagcc attgtcgagt cctccaaaca
 1441 accgtatcgt tgtgtcacct ggtaaagtga tgtcaccgtt gcaggcgagg actccttccc
 1501 cgcaatacca gcagtcgttt cttgttcagg gcgcttcggg cgtaagcaaa acttacgtat
 1561 caatgggcca gagttccgga ggacaacttc ttgtgccttt tatgctcgat cagcaacaac
 1621 agcagcagca gcaaaccgcc attacaaatg ccaacgagat cggaagattt ttattcaata
 1681 tggacaccca gcaaactagt gtggaagtca accaatatga agtaattgat cattccttgt
 1741 cgaacaacct gtctgcgagt cttaatctta tagaacagga ttaccaaccg gacgctagct
 1801 cggaaagcct gacgagactt actaccaatg cgtacaatca attaaccaac aattttccat
 1861 agaaaacaaa acatattatc tcatcggcat aatattgttt aaattcaaat aacgctgaga
 1921 ggccatctat ttaattaaga gatctcaaac aaaatgatga atttatttgt aaatatctat
 1981 aaatttgtga taagattatt tttcaaaaaa aaaaaaa
//
```

```
LOCUS       Drosocin, isoform A        1571 bp    DNA     linear   UNA 03-MAR-2026
DEFINITION  Pyrrhocoris apterus Drosocin, isoform A, complete mRNA.
ACCESSION
KEYWORDS    .
SOURCE      Pyrrhocoris apterus
  ORGANISM  Pyrrhocoris apterus
            cellular organisms; Eukaryota; Opisthokonta; Metazoa; Eumetazoa;
            Bilateria; Protostomia; Ecdysozoa; Panarthropoda; Arthropoda;
            Mandibulata; Pancrustacea; Hexapoda; Insecta; Dicondylia;
            Pterygota; Neoptera; Paraneoptera; Hemiptera; Prosorrhyncha;
            Heteroptera; Euheteroptera; Neoheteroptera; Panheteroptera;
            Pentatomomorpha; Pyrrhocoroidea; Pyrrhocoridae; Pyrrhocoris.
FEATURES             Location/Qualifiers
     5'UTR           1..48
                     /gene="Dro"
     gene            1..1571
                     /gene="Dro"
     mRNA            join(1..26,27..192,193..333,334..474,475..615,616..756,
                     757..897,898..1038,1039..1179,1180..1571)
                     /gene="Dro"
                     /note="Drosocin, isoform A"
     source          1..>1571
                     /mol_type="mRNA"
                     /organism="Pyrrhocoris apterus"
                     /db_xref="taxon:37000"
     CDS             join(49..192,193..333,334..474,475..615,616..756,757..897,
                     898..1038,1039..1179,1180..1305)
                     /gene="Dro"
                     /product="Drosocin, isoform A"
                     /translation="MVKVSTYGICILSIIGLAMVEAVDKGGYLPRPTPPRQIYNRNKR
                     EVGDEPFADLAPESRQVRSVDFISAVDKGSYLPRPTPPRPIYNRNKREVGDEPFADLA
                     PESRQVRSVDFISAVDKGSYLPRPTPPRPIYNRNKREVGDEPFADLAPESRQVRSVDF
                     ISAVDKGSYLPRPTPPRPIYNRNKREVGDEPFADLAPENRQVRSVDFISAVDKGSYLP
                     RPTPPRPIYNRNKREVGDEPFADLAPESRQVRSVDFISAVDKGSYLPRPTPPRPIYNR
                     NKREVGDEPFADLAPENRQVRSVDFISAVDKGSYLPRPTPPRPIYNRNKREVGDEPFA
                     DLAPESRQVRSVDFISAVDKGSYLPRPTPPRPIYNRNKREVGDESFADLAPENRQVRS
                     VDFISAVDKGSYSLGLLLRGRSFPDF"
                     /codon_start=1
     primer_bind     1277..1296
                     /standard_name="Dro-Fwd"
     3'UTR           1306..1571
                     /gene="Dro"
     primer_bind     complement(1415..1434)
                     /standard_name="Dro-Rev"
ORIGIN
        1 tcagaatagt cctgcagttg gaaccgaata atctgacata ctgttataat ggtgaaagta
       61 agcacttacg gcatatgcat cctttccatc atcgggctag caatggtaga agcagttgac
      121 aaaggtggct atctccctag gcctactcct ccaaggcaga tctacaacag aaataagaga
      181 gaggtcggtg atgaaccctt cgcagaccta gcaccggaaa gccgtcaagt tagaagtgtt
      241 gactttataa gtgcagttga caagggtagc tatctcccta ggcctactcc tccaaggccg
      301 atctacaaca gaaataagag agaggtcggt gatgaaccct tcgcagacct agcaccggaa
      361 agccgtcaag ttagaagtgt tgactttata agtgcagttg acaagggtag ctatctccct
      421 aggcctactc ctccaaggcc gatctacaac agaaataaga gagaggtcgg tgatgaaccc
      481 ttcgcagacc tagcaccgga aagccgtcaa gttagaagtg ttgactttat aagtgcagtt
      541 gacaagggta gctatctccc taggcctact cctccaaggc cgatctacaa cagaaataag
      601 agagaggtcg gtgatgaacc cttcgcagac ctagcaccgg aaaaccgtca agttagaagt
      661 gttgacttta taagtgcagt tgacaagggt agctatctcc ctaggcctac tcctccaagg
      721 ccgatctaca acagaaataa gagagaggtc ggtgatgaac ccttcgcaga cctagcaccg
      781 gaaagccgtc aagttagaag tgttgacttt ataagtgcag ttgacaaggg tagctatctc
      841 cctaggccta ctcctccaag gccgatctac aacagaaata agagagaggt cggtgatgaa
      901 cccttcgcag acctagcacc ggaaaaccgt caagttagaa gtgttgactt tataagtgca
      961 gttgacaagg gtagctatct ccctaggcct actcctccaa ggccgatcta caacagaaat
     1021 aagagagagg tcggtgatga acccttcgca gacctagcac cggaaagccg tcaagttaga
```

```
1081 agtgtagact ttataagtgc agttgacaag ggtagctatc tccctaggcc tactcctcca
1141 aggccgatct acaacagaaa taagagagag gtcggtgatg aatccttcgc agacctagca
1201 ccggaaaacc gtcaagttag aagtgttgac tttataagtg cagttgacaa gggtagctac
1261 tccctaggcc tactcctacg aggccgatct tttcctgatt tttaaaaatt ttcaattttt
1321 ttactcatat ttctgaatag gcgaggatat atttttcgtt ttctaaaatt gaaatttaaa
1381 ctgatttgtt ccgtttcgaa ctgccaatgt ttgctggtgc tgctggttaa atgcaaatat
1441 tgtttgtgat atgtttatac tttaaattta cttttatatt atagttttta ataatcatac
1501 agaaacgttt tatatgcata tttcagtgtt gtcaaagaac caatattaat aaactgtatt
1561 gtattgttat a
//
```

```
LOCUS       Hemiptericin-1           962 bp    DNA     linear   UNA 03-MAR-2026
DEFINITION  Pyrrhocoris apterus Hemiptericin-1, complete mRNA.
ACCESSION
KEYWORDS    .
SOURCE      Pyrrhocoris apterus
  ORGANISM  Pyrrhocoris apterus
            cellular organisms; Eukaryota; Opisthokonta; Metazoa; Eumetazoa;
            Bilateria; Protostomia; Ecdysozoa; Panarthropoda; Arthropoda;
            Mandibulata; Pancrustacea; Hexapoda; Insecta; Dicondylia;
            Pterygota; Neoptera; Paraneoptera; Hemiptera; Prosorrhyncha;
            Heteroptera; Euheteroptera; Neoheteroptera; Panheteroptera;
            Pentatomomorpha; Pyrrhocoroidea; Pyrrhocoridae; Pyrrhocoris.
FEATURES             Location/Qualifiers
     5'UTR           1..48
                     /gene="Hmp1"
     gene            1..962
                     /gene="Hmp1"
     mRNA            join(1..40,41..171,172..962)
                     /gene="Hmp1"
                     /note="Hemiptericin-1"
     source          1..962
                     /organism="Pyrrhocoris apterus"
                     /mol_type="mRNA"
                     /db_xref="taxon:37000"
                     /note="contig_1016_segment0"
     CDS             join(49..171,172..597)
                     /gene="Hmp1"
                     /product="Hemiptericin-1"
                     /translation="MRFQSSLIFLCIAVFASALGAFIEIPDEYLQDSAALQFDGEDEL
                     WRSKRDVELKGKGGENEGFVGLKAQRNLYEDDRTSLSGTVKGQSQWKDPYPAQHAGMA
                     RLDGTRTLIENDRTKVTGSGFAQREVATGMRPHDSFGVGVEATHNIYKGKNGEVDVFG
                     GVQRQWNTPDRHQARGGIRWRF"
                     /codon_start=1
     primer_bind     274..291
                     /standard_name="Hmp1-Fwd"
     primer_bind     complement(502..520)
                     /standard_name="Hmp1-Rev"
     3'UTR           598..962
                     /gene="Hmp1"
ORIGIN
        1 agtccgtcat cagtcttaca gtagagatct aataacaaag gagcaaacat gcgattccaa
       61 tcgtcactca ttttcctctg tatcgctgtg ttcgcatcag cgttgggggc cttcatcgaa
      121 atacctgatg aatatctaca agattctgct gctctacaat tcgatggaga ggacgaactc
      181 tggaggtcaa aacgagacgt agaactgaaa ggtaaaggag gagaaaatga aggattcgtc
      241 gggttgaaag cccagcgtaa tctctatgag gacgacagga ccagcctcag tggtacggta
      301 aagggccaga gccagtggaa ggatccctat cctgcacagc acgctggaat ggctaggctt
      361 gacggaacac gaaccctgat cgagaatgac aggaccaaag ttacgggatc agggttcgcc
      421 cagagggaag tggccactgg aatgcgtcct catgactctt tcggtgttgg agtcgaagcc
      481 acacacaaca tatacaaggg gaagaacggc gaggtggatg tgttcggagg agtacagagg
      541 cagtggaaca ctccagacag acatcaagcc aggggtggaa tcaggtggag attttaggag
      601 gacatagtaa accaacaaaa ccaatttgta ataatgtaaa ataaatgttt aagataatgt
      661 ataaatttgt atattgtgta aagtgtaaaa aagaaaaaaa atatatattt acaaagaaat
      721 ggaaataaat aatgaattaa atggcttcaa cattcttgag gaatttctat attaaaatgt
      781 atctttatag aaaaataatt gttttttttt ttaagcgaca ttcagtgtgt cggcggtatc
      841 gaaacaccag tacaaatgta aatattaaag ttttgtgtaa cgttaacaag ttgaattatg
      901 tcaaatatca accaacaaac atgtatcttt attgaaaaat aataatataa taatattaat
      961 aa
//


LOCUS       Hemiptericin-2           935 bp    DNA     linear   UNA 03-MAR-2026
DEFINITION  Pyrrhocoris apterus Hemiptericin-2, complete mRNA.
ACCESSION
```

```
KEYWORDS    .
SOURCE      Pyrrhocoris apterus
  ORGANISM  Pyrrhocoris apterus
            cellular organisms; Eukaryota; Opisthokonta; Metazoa; Eumetazoa;
            Bilateria; Protostomia; Ecdysozoa; Panarthropoda; Arthropoda;
            Mandibulata; Pancrustacea; Hexapoda; Insecta; Dicondylia;
            Pterygota; Neoptera; Paraneoptera; Hemiptera; Prosorrhyncha;
            Heteroptera; Euheteroptera; Neoheteroptera; Panheteroptera;
            Pentatomomorpha; Pyrrhocoroidea; Pyrrhocoridae; Pyrrhocoris.
FEATURES             Location/Qualifiers
     5'UTR           1..40
                     /gene="Hmp2"
     gene            1..935
                     /gene="Hmp2"
     mRNA            join(1..32,33..163,164..935)
                     /gene="Hmp2"
                     /note="Hemiptericin-2"
     source          1..935
                     /organism="Pyrrhocoris apterus"
                     /mol_type="mRNA"
                     /db_xref="taxon:37000"
                     /note="contig_1016_segment0"
     CDS             join(41..163,164..589)
                     /gene="Hmp2"
                     /product="Hemiptericin-2"
                     /translation="MRFQSSLIFLGVAVFASALGAFVEIPDEYLQDSAALQFDGEDEL
                     WRSKRDVELKGKGGENEGFVGLKAQRNLYEDDRTSLSGTVKGQSQWKDPYPAQHAGMA
                     RLDGTRTLIENDRTKVTGSGFAQREVATGMRPHDSFGVGVEATHNIYKGKNGEVDVFG
                     GVQRQWNTPDRHQARGGIRWRF"
                     /codon_start=1
     primer_bind     266..283
                     /standard_name="Hmp2-Fwd"
     primer_bind     complement(494..512)
                     /standard_name="Hmp2-Rev"
     3'UTR           590..935
                     /gene="Hmp2"
ORIGIN
        1 atcagtctta cagtagagac ctaataacaa aggtgtgaaa atgcgattcc aatcctcact
       61 tattttcctc ggcgtcgctg tgtttgcatc agcgttgggg gcctttgtcg aaataccaga
      121 tgaatatcta caagattctg cagccctaca attcgatgga gaggacgaac tctggaggtc
      181 aaaacgagac gtagaactga aaggtaaagg aggagaaaat gaaggattcg tcgggttgaa
      241 agcccagcgt aatctctatg aggacgacag gaccagcctc agtggtacgg taaagggcca
      301 gagccagtgg aaggatccct atcctgcaca gcacgctgga atggctaggc ttgacggaac
      361 acgaaccctg atcgagaatg acaggaccaa agttacggga tcagggttcg cccagaggga
      421 agtggccact ggaatgcgtc ctcatgactc tttcggtgtt ggagtcgaag ccacacacaa
      481 catatacaag gggaagaacg gcgaggtgga tgtgttcgga ggagtacaga ggcagtggaa
      541 cactccagac agacatcaag ccaggggtgg aatcaggtgg agattttagg aggacatagt
      601 aaaccaacaa aaccaatttg taataatgta caataaatgt tttagataat gtataaattt
      661 gtatattgtg taaagtgtaa aaaagaaaaa aaaaatatat atttacaaag aaaaggaaat
      721 aaataatgaa ttaaatggct tcaacattct tgaggaattt ctatattaaa atgtatcttt
      781 atagaaaaat aattgttttt ttttaagcga cattcagtgt gtcggcggta tcgaaacacc
      841 agtacaaatg taaatattaa agttttgtgt aacgttaaca agttgaatta tgtcaaatat
      901 caaccaacaa acatgtatct ttattgaaaa ataat
//
```

```
LOCUS       Hemiptericin-3           844 bp    DNA     linear   UNA 03-MAR-2026
DEFINITION  Pyrrhocoris apterus Hemiptericin-3, complete mRNA.
ACCESSION
KEYWORDS    .
SOURCE      Pyrrhocoris apterus
  ORGANISM  Pyrrhocoris apterus
            cellular organisms; Eukaryota; Opisthokonta; Metazoa; Eumetazoa;
            Bilateria; Protostomia; Ecdysozoa; Panarthropoda; Arthropoda;
            Mandibulata; Pancrustacea; Hexapoda; Insecta; Dicondylia;
            Pterygota; Neoptera; Paraneoptera; Hemiptera; Prosorrhyncha;
            Heteroptera; Euheteroptera; Neoheteroptera; Panheteroptera;
            Pentatomomorpha; Pyrrhocoroidea; Pyrrhocoridae; Pyrrhocoris.
FEATURES             Location/Qualifiers
     5'UTR           1..42
                     /gene="Hmp3"
     gene            1..844
                     /gene="Hmp3"
     mRNA            join(1..34,35..165,166..844)
                     /gene="Hmp3"
                     /note="Hemiptericin-3"
     source          1..844
                     /organism="Pyrrhocoris apterus"
                     /mol_type="mRNA"
                     /db_xref="taxon:37000"
                     /note="contig_1016_segment0"
     CDS             join(43..165,166..591)
                     /gene="Hmp3"
                     /product="Hemiptericin-3"
                     /translation="MRFQSSLIFLCIAVFASGLGAFIEIPDEYLQDSAALQFDGEDEL
                     WRSKRDVELKGKGGENEGFVGLKAQRILYEDDRTSLSGTVKGQSQWKDPYPAQHAGMA
                     RLDGTRTLFENDRTKVTGSGFAQREVATGMRPHDSFGVGVEATHNIYKGKNGEVDVFG
                     GVQRQWNTPDRHQARGGIRWRF"
                     /codon_start=1
     primer_bind     268..285
                     /standard_name="Hmp3-Fwd"
     primer_bind     complement(496..514)
                     /standard_name="Hmp3-Rev"
     3'UTR           592..844
                     /gene="Hmp3"
ORIGIN
        1 tcatcagtct tacagtagag atctaataac aaaggagcaa acatgcgatt ccaatcgtca
       61 ctcattttcc tctgtatcgc tgtgttcgcg tcagggttgg gggccttcat cgaaatacct
      121 gatgaatatc tacaagattc tgctgctcta caattcgatg gagaggacga actctggagg
      181 tcaaaacggg acgtagaact gaaaggtaaa ggaggagaaa atgaaggatt cgtcgggttg
      241 aaagcccagc gtatactcta tgaggacgac aggaccagcc tcagtggtac agtaaagggc
      301 cagagccagt ggaaggatcc ctatcctgca cagcacgctg gaatggctag gcttgacgga
      361 acacgaaccc tgttcgagaa tgacaggacc aaagttacgg gatcagggtt cgcccagagg
      421 gaagtggcca ctggaatgcg tcctcatgac tctttcggtg ttggagtcga agccacacac
      481 aacatataca aggggaagaa cggcgaggtg gatgtgttcg gaggagtaca gaggcagtgg
      541 aacaccccag acagacatca agccaggggt ggaatcaggt ggagatttta ggaggactta
      601 gtaaaccaac aaaaccaatt tgttaataat gtacaatact ttttttagtt catctataaa
      661 tttgtatatt gtgtaaagtg taaaaaaaag aagaaggaaa tgtgtattta cacaggaaag
      721 gaaattatca atacaaatgt aatatattac agtttttttt atgacattac aaatttgcat
      781 tatgtacaat taagaccaat aaacatgtat atttattgtg taataaattt ttaatttaat
      841 gaca
//
LOCUS       Hemiptericin-4           922 bp    DNA     linear   UNA 03-MAR-2026
DEFINITION  Pyrrhocoris apterus Hemiptericin-4, complete mRNA.
ACCESSION
KEYWORDS    .
SOURCE      Pyrrhocoris apterus
  ORGANISM  Pyrrhocoris apterus
            cellular organisms; Eukaryota; Opisthokonta; Metazoa; Eumetazoa;
```

```
            Bilateria; Protostomia; Ecdysozoa; Panarthropoda; Arthropoda;
            Mandibulata; Pancrustacea; Hexapoda; Insecta; Dicondylia;
            Pterygota; Neoptera; Paraneoptera; Hemiptera; Prosorrhyncha;
            Heteroptera; Euheteroptera; Neoheteroptera; Panheteroptera;
            Pentatomomorpha; Pyrrhocoroidea; Pyrrhocoridae; Pyrrhocoris.
FEATURES             Location/Qualifiers
     5'UTR           1..10
                     /gene="Hmp4"
     gene            1..922
                     /gene="Hmp4"
     mRNA            join(1..133,134..922)
                     /gene="Hmp4"
                     /note="Hemiptericin-4"
     source          1..922
                     /organism="Pyrrhocoris apterus"
                     /mol_type="mRNA"
                     /db_xref="taxon:37000"
                     /note="contig_1016_segment0"
     CDS             join(11..133,134..559)
                     /gene="Hmp4"
                     /product="Hemiptericin-4"
                     /translation="MRFQSSLIFLCIAVFASALGAFIEIPDEYLQDSAALQFDGEDEL
                     WRSKRDVELKGKGGENEGFVGLKAQRILYEDDRTSLSGTVKGQSQWKDPYPAQHAGMA
                     RLDGTRTLIENDRTKVTGSGFAQREVATGMRPHDSFGVGVEATHNIYKGKNGEVDVFG
                     GVQREWNTPDRHQARGGIRWRF"
                     /codon_start=1
     primer_bind     236..253
                     /standard_name="Hmp4-Fwd"
     primer_bind     complement(464..482)
                     /standard_name="Hmp4-Rev"
     3'UTR           560..922
                     /gene="Hmp4"
ORIGIN
        1 aggagcaaac atgcgattcc aatcgtcact cattttcctc tgtatcgctg tgttcgcatc
       61 agcgttgggg gccttcatcg aaatacctga tgaatatcta caagattctg ctgctctaca
      121 attcgatgga gaggatgaac tctggaggtc aaaacgagac gtagaactga aaggtaaagg
      181 aggagaaaat gaaggattcg tcgggttgaa agcccagcgt atactctatg aggacgacag
      241 gaccagcctc agtggtacag taaagggcca gagccagtgg aaggatccct atcctgcaca
      301 gcacgctgga atggctaggc ttgacggaac ccgaaccctg atcgagaatg acaggaccaa
      361 agttacggga tcagggttcg cccagaggga agtggccact ggaatgcgtc ctcatgactc
      421 tttcggtgtt ggagtcgaag ccacacacaa catatacaag ggaagaacg gcgaggtgga
      481 tgtgttcgga ggagtacaga gagagtggaa cactccagac agacatcaag ccaggggtgg
      541 aatcaggtgg agattttagg aggacatagt aaaccaacaa aaccaatttg taataatgta
      601 aaataaatgt ttaagataat gtataaattt gtatattgtg taaagtgtaa aaaagaaaaa
      661 aaatatatat ttacaaagaa aaggaaataa ataatgaatt aaatggcttc aacattcttg
      721 aggaatttct atattaaaat gtatctttat agaaaaataa ttgttttttt ttaagcgaca
      781 ttcagtgtgt cggcggtatc gaaacaccag tacaaatgta aatattaaag ttttgtgtaa
      841 cgttaacaag ttgaattatg tcaaatatca accaacaaac atgtatcttt attgaaaaat
      901 aataatataa taatattaat aa
//
```

```
LOCUS       Hemiptericin-5           939 bp    DNA     linear   UNA 03-MAR-2026
DEFINITION  Pyrrhocoris apterus Hemiptericin-5, complete mRNA.
ACCESSION
KEYWORDS    .
SOURCE      Pyrrhocoris apterus
  ORGANISM  Pyrrhocoris apterus
            cellular organisms; Eukaryota; Opisthokonta; Metazoa; Eumetazoa;
            Bilateria; Protostomia; Ecdysozoa; Panarthropoda; Arthropoda;
            Mandibulata; Pancrustacea; Hexapoda; Insecta; Dicondylia;
            Pterygota; Neoptera; Paraneoptera; Hemiptera; Prosorrhyncha;
            Heteroptera; Euheteroptera; Neoheteroptera; Panheteroptera;
            Pentatomomorpha; Pyrrhocoroidea; Pyrrhocoridae; Pyrrhocoris.
FEATURES             Location/Qualifiers
     5'UTR           1..41
                     /gene="Hmp5"
     gene            1..939
                     /gene="Hmp5"
     mRNA            join(1..33,34..164,165..939)
                     /gene="Hmp5"
                     /note="Hemiptericin-5"
     source          1..939
                     /organism="Pyrrhocoris apterus"
                     /mol_type="mRNA"
                     /db_xref="taxon:37000"
                     /note="contig_2129_segment1"
     CDS             join(42..164,165..590)
                     /gene="Hmp5"
                     /product="Hemiptericin-5"
                     /translation="MRFQSSLIFLGVAVFASALGAFIEIPDEYLQDSAALQFDGEDEL
                     WRSKRDVELKGKGGENEGFVGLKAQRNLYEDDRTSLSGTVKGQSQWKDHYPAQHAGMA
                     RLDGTRTLFENDRTKVTGSGFAQREVATGMRPHDSFGVGVEATHNIYKGKNGEVDVFG
                     GVQREWNTPDRHQARGGIRWRF"
                     /codon_start=1
     primer_bind     267..284
                     /standard_name="Hmp5-Fwd"
     primer_bind     complement(495..513)
                     /standard_name="Hmp5-Rev"
     3'UTR           591..939
                     /gene="Hmp5"
ORIGIN
        1 catcagtctt acagtagaga cctaataaca aaggtgtgaa aatgcgattc caatcctcac
       61 ttattttcct cggcgtcgct gtgtttgcat cagcgttggg ggccttcatc gaaatacctg
      121 atgaatatct acaagattct gcagctctac aattcgatgg agaggacgaa ctctggaggt
      181 caaaacggga cgtagaactg aaaggtaaag gaggagaaaa tgaaggattc gtcgggttga
      241 aagcccagcg taatctctat gaggacgaca ggaccagcct cagtggtaca gtaaagggcc
      301 agagccagtg gaaggatcat tatcctgcac agcacgctgg aatggccagg ctcgacggaa
      361 cacgaaccct gttcgagaat gacaggacca aagtcacggg atcagggttc gcccagaggg
      421 aagtggccac tggaatgcgt cctcatgact ctttcggtgt tggagtcgaa gccacacaca
      481 acatatacaa ggggaagaac ggcgaggtgg atgtgttcgg aggagtacag agagagtgga
      541 acactccaga cagacatcaa gccaggggtg gaatcaggtg gagattttag gaggatatag
      601 taaaccaaga aaaccaattt gtaataatgt acaataatat ttttagataa tgtataaatt
      661 tgtataatgt gtaaagtgta aaaaaggaaa aaattatata tttacaaaga aaaagaaaga
      721 aataatgaat taaatggctt aatcattctt gagcactttc tatattaaaa tgtatcttta
      781 tagaaaaata attgtttgtt tttttaagtg acaccttcag tgtgtcccac tgtgatacca
      841 gtacaaatgt aaatattaaa gttttgtata acgttaacaa gttgaattat atcaaatatc
      901 aaccaacaaa catgtatctt taaagaaaaa taattgttt
//
LOCUS       Hemiptericin-6           931 bp    DNA     linear   UNA 03-MAR-2026
DEFINITION  Pyrrhocoris apterus Hemiptericin-6, complete mRNA.
ACCESSION
KEYWORDS    .
SOURCE      Pyrrhocoris apterus
  ORGANISM  Pyrrhocoris apterus
```

```
            cellular organisms; Eukaryota; Opisthokonta; Metazoa; Eumetazoa;
            Bilateria; Protostomia; Ecdysozoa; Panarthropoda; Arthropoda;
            Mandibulata; Pancrustacea; Hexapoda; Insecta; Dicondylia;
            Pterygota; Neoptera; Paraneoptera; Hemiptera; Prosorrhyncha;
            Heteroptera; Euheteroptera; Neoheteroptera; Panheteroptera;
            Pentatomomorpha; Pyrrhocoroidea; Pyrrhocoridae; Pyrrhocoris.
FEATURES             Location/Qualifiers
     5'UTR           1..40
                     /gene="Hmp6"
     gene            1..931
                     /gene="Hmp6"
     mRNA            join(1..32,33..163,164..931)
                     /gene="Hmp6"
                     /note="Hemiptericin-6"
     source          1..931
                     /organism="Pyrrhocoris apterus"
                     /mol_type="mRNA"
                     /db_xref="taxon:37000"
                     /note="contig_2129_segment1"
     CDS             join(41..163,164..589)
                     /gene="Hmp6"
                     /product="Hemiptericin-6"
                     /translation="MRFQSSLIFLCIAVFASALGAFIEIPDEYLQDSAALQFDGEDEL
                     WRSKRDVELKGKGGENEGFVGLKAQRNLYEDDRTSLSGTVKGQSQWKDPYPAQHAGMA
                     RLDGTRTLFENDRTKVTGSGFAQREVATGMRPHDSFGAGVEATHNIYKGKNGEVDVFG
                     GVQREWNTPDRHQARGGIRWRF"
                     /codon_start=1
     primer_bind     266..283
                     /standard_name="Hmp6-Fwd"
     primer_bind     complement(494..512)
                     /standard_name="Hmp6-Rev"
     3'UTR           590..931
                     /gene="Hmp6"
ORIGIN
        1 atcagtctta cagtagagac ctaataacaa aggggcaaac atgcgattcc aatcgtcact
       61 cattttcctc tgtatcgctg tgttcgcgtc agcgttgggg gccttcatcg aaatacctga
      121 tgaatatcta caagattctg ctgctctaca attcgacgga gaggatgaac tctggaggtc
      181 aaaacgagac gtagaactga aaggtaaagg aggagaaaat gaaggattcg tcgggttgaa
      241 agcccagcgt aatctctatg aggacgacag gaccagcctc agtggtacag taaagggcca
      301 gagccagtgg aaggatccct atcctgcaca gcacgctgga atggccaggc tcgacggaac
      361 acgaaccctg ttcgagaatg acaggaccaa agttacggga tcagggttcg cccagaggga
      421 agtggccact ggaatgcgtc ctcatgactc tttcggtgct ggagtcgaag ccacacacaa
      481 catttacaag ggaaagaacg gcgaggtgga tgtgttcgga ggagtacaga gagagtggaa
      541 cactccagac agacatcaag ccaggggtgg aatcaggtgg agattttagg aggacttagt
      601 aaatcaagaa aaccaatttg taataatgta caataatatt tttagataat gtataaattt
      661 gtataatgtg taaagtgtaa aaaaaaaaaa attatatatt tacaaataaa aagaaagaaa
      721 taatgaatta aatggcttca tcattcttga gcactttcta tattaaaatg tatctttata
      781 gaaaaataat tgtttttttt tttaagtgac accttcagtg tgtcccactg tgataccagt
      841 acaaatgtaa atattaaagg tttgaataac gttaacaagt tgaattatgt caaatatcaa
      901 ccaacaaaca tgtatcttta aagaaaaata a
//
```

```
LOCUS       Hemiptericin-7          820 bp    DNA     linear   UNA 03-MAR-2026
DEFINITION  Pyrrhocoris apterus Hemiptericin-7, complete mRNA.
ACCESSION
KEYWORDS    .
SOURCE      Pyrrhocoris apterus
  ORGANISM  Pyrrhocoris apterus
            cellular organisms; Eukaryota; Opisthokonta; Metazoa; Eumetazoa;
            Bilateria; Protostomia; Ecdysozoa; Panarthropoda; Arthropoda;
            Mandibulata; Pancrustacea; Hexapoda; Insecta; Dicondylia;
            Pterygota; Neoptera; Paraneoptera; Hemiptera; Prosorrhyncha;
            Heteroptera; Euheteroptera; Neoheteroptera; Panheteroptera;
            Pentatomomorpha; Pyrrhocoroidea; Pyrrhocoridae; Pyrrhocoris.
FEATURES             Location/Qualifiers
     5'UTR           1..40
                     /gene="Hmp7"
     gene            1..820
                     /gene="Hmp7"
     mRNA            join(1..32,33..163,164..820)
                     /gene="Hmp7"
                     /note="Hemiptericin-7"
     source          1..820
                     /organism="Pyrrhocoris apterus"
                     /mol_type="mRNA"
                     /db_xref="taxon:37000"
                     /note="contig_2129_segment1"
     CDS             join(41..163,164..589)
                     /gene="Hmp7"
                     /product="Hemiptericin-7"
                     /translation="MRFQSSLIFLCIAVFASALGAFIEIPDEYLQDSAALQFDGEDEL
                     WRSKRDVELKGKGGENEGFVGLKAQRNLYEDDRTSLSGTVKGQSQWKDPYPAQHAGMA
                     RLDGTRTLFENDRTKVTGSGFAQREVATGMRPHDSFGVGVEATHNIYKGKNGEVDVFG
                     GVQRQWNTPDRHQARGGIRWRF"
                     /codon_start=1
     primer_bind     266..283
                     /standard_name="Hmp7-Fwd"
     primer_bind     complement(494..512)
                     /standard_name="Hmp7-Rev"
     3'UTR           590..820
                     /gene="Hmp7"
ORIGIN
        1 atcagtctta cagtagagac ctaataacaa aggggcaaac atgcgattcc aatcgtcact
       61 cattttcctc tgtatcgctg tgttcgcgtc agcgttgggg gccttcatcg aaatacctga
      121 tgaatatcta caagattctg ctgctctaca attcgacgga gaggacgaac tctggaggtc
      181 aaaacgagac gtagaactga aaggtaaagg aggagaaaat gaaggattcg tcgggttgaa
      241 agcccagcgt aatctctatg aggacgacag gaccagcctc agtggtacgg taaagggcca
      301 gagccagtgg aaggatccct atcctgcaca gcacgctgga atggctaggc tcgacggaac
      361 acgaaccctg ttcgagaatg acaggaccaa agttacggga tcagggttcg cccagaggga
      421 agtggccact ggaatgcgtc ctcatgactc tttcggtgtt ggagtcgaag ccacacacaa
      481 catttacaag ggaaagaacg gcgaggtgga tgtgttcgga ggagtacaga ggcagtggaa
      541 caccccagac agacatcaag ccaggggtgg aatcaggtgg agattttagg aggacttagt
      601 aaaccaacaa aaccaatttg ttaataatgt acaatacttt ttttagttca tctataaatt
      661 tgtatattgt gtaaagtgta aaaaaaagaa ggaaatgtgt atttacacag aaaaggaaac
      721 tatcaataca aatgtaatat attacagttt ttatgacatt aagaatttac attatgtaca
      781 attaagacta atagacatgt atatttatga tgtaataaaa
//
LOCUS       Peroxidasin            4575 bp    DNA     linear   UNA 03-MAR-2026
DEFINITION  Pyrrhocoris apterus Peroxidasin, complete mRNA.
ACCESSION
KEYWORDS    .
SOURCE      Pyrrhocoris apterus
  ORGANISM  Pyrrhocoris apterus
            cellular organisms; Eukaryota; Opisthokonta; Metazoa; Eumetazoa;
            Bilateria; Protostomia; Ecdysozoa; Panarthropoda; Arthropoda;
```

```
            Mandibulata; Pancrustacea; Hexapoda; Insecta; Dicondylia;
            Pterygota; Neoptera; Paraneoptera; Hemiptera; Prosorrhyncha;
            Heteroptera; Euheteroptera; Neoheteroptera; Panheteroptera;
            Pentatomomorpha; Pyrrhocoroidea; Pyrrhocoridae; Pyrrhocoris.
FEATURES             Location/Qualifiers
     5'UTR           1..61
                     /gene="Pxn"
     gene            1..4575
                     /gene="Pxn"
     mRNA            join(1..231,232..303,304..375,376..447,448..519,520..591,
                     592..743,744..861,862..1028,1029..1277,1278..1439,
                     1440..1710,1711..1832,1833..1977,1978..2090,2091..2321,
                     2322..2563,2564..2719,2720..2920,2921..3104,3105..3256,
                     3257..3602,3603..3720,3721..3929,3930..4153,4154..4575)
                     /gene="Pxn"
                     /note="Peroxidasin"
     source          1..4575
                     /mol_type="mRNA"
                     /organism="Pyrrhocoris apterus"
                     /db_xref="taxon:37000"
     CDS             join(62..231,232..303,304..375,376..447,448..519,520..591,
                     592..743,744..861,862..1028,1029..1277,1278..1439,
                     1440..1710,1711..1832,1833..1977,1978..2090,2091..2321,
                     2322..2563,2564..2719,2720..2920,2921..3104,3105..3256,
                     3257..3602,3603..3720,3721..3929,3930..4153,4154..4249)
                     /gene="Pxn"
                     /product="Peroxidasin"
                     /translation="MTGLSVLLLIATVLGLVALSTATTVCPRRCVCIRTTVRCMHLRL
                     DYIPSAPVNTTILDLRFNRIRQIPPGTFSHLKNLDTLMLSDNLIQSLENGLFTGLEQL
                     RHLYLYKNKIRNIEEGAFDGMPKLEQLYLHYNKIMKIHPNLFNNLKNLQRLFLHENRI
                     HYIPRGTFANMAYLKRVRLDGNALVCDCGLKEMMMAMPKAEVAAVCQYPENMRGKTLH
                     EMTPSDIVCNEPRIIEGPHDVEVSFGGSAIFTCRVEADSDAEIIWMLNSNEIDKSDPR
                     YSVSQDGTLVVKSMTDKEMGVYECMVKSTNGMSKSQAAKTLYNHVSGKAEWKRKPKDQ
                     SAREGDTVQLECAVSNGHVTWTRNGVPISAPVSAEGTLTLSSVTPQDTASYTCNVETP
                     QGKIKAEAKLTVSAPPRILQPPESSSVRSGETVLLRCRASGTPEPILSWFKDGASLQP
                     SERIIYNDDFSEVTLKKVSLGDSGLYTCMAQSTIGTAEESGELRVRPVGPRPPAFLIA
                     PYPIIAQAGTSVELPCKVQGDPYPTIEWLKNGEPIQLDQTHRVFPIGSLRLYNITQSD
                     AGIYRCTATNVHGEISAHATLTVEGANNIDDSFVLTAIQNAKDSVDRAVNSTLDSIFS
                     KKEPKNPSALMRLLRYPDEQARSVVRAADIYETTLLNIRKHIISGLQFNETEDFHYEE
                     VLSPDQLQLIARMSGCNEHQKANCSNMCFHTKYRTIDGTCNNLQHPYWGASHTAFRRL
                     LKPIYENGFSQPVGWDKNRKYNGFSLPNARKVSLTLIRTENITHDDTITHMVMQWGQF
                     LDHDLDHAIPSTTAESWEGVDCKKTCRATPPCFPMEVEQDDPRVKNRRCMDFIRSSAI
                     CGSGMTSVFFNTIQPREQINQLTSFIDATQVYGFHDNRSILLRDYTEDLGLLRVGLIT
                     DSGKPLLPVAGAQEVDCRRDPTENEAGCFLAGDIRVNEQISLVAMHTIWLREHNRIAT
                     ELKEINPHWTGEIIFQEARKIVGAQMQHITYKHWIRNILGEKGMQILGEYEGYDPNLD
                     AGISNVFATAALRFGHTLINPVLARLDSEFKTIPEGDLPLGRAFFAPWRIIEEGGVDA
                     LMRGMWMSAAKKKMPNQNLNNELTDHLFTSFHAVALDLASMNVQRSRDHGIPFYNEFR
                     KLCNLSSVQTFDDLKEQISNFEVREKLKNLYGHPHNIDIFVGGILEDQIDGAKFGPTF
                     RCLLIEQFKRIRNGDRFWYENPATFKASQLTQIRQSSLARVLCDNGDNITRISPDVFI
                     LPEKQTPSIVDCSDIPKIDLRFWYECEDCENDVSSRGRRGISGEDMEERMEGLENMVQ
                     QLQKTVKSLKKKVNGLSKCRDNKGVNRKDGDTWQKDNCTLCECQGKQVSCTRIQCAKL
                     TCQRQEKVEGRCCPVCV"
                     /codon_start=1
     primer_bind     2581..2602
                     /standard_name="Pxd-Fwd"
     primer_bind     complement(2715..2735)
                     /standard_name="Pxd-Rev"
     3'UTR           4250..4575
                     /gene="Pxn"
ORIGIN
        1 atagtagcgg gcggcctgtc gtgtggcgag ggttagtgct gttcagtagt gttcgggaga
       61 gatgacggga ctgtctgtct tactactgat agccacagtc ctaggactgg tggcgttatc
      121 aacagcaaca accgtatgtc cgaggaggtg cgtctgtata aggactacag tgcgatgcat
```

181 gcacctgcgt ctagattata taccatcggc tcctgtcaac acaacaatac tggacttgag
241 gttcaacaga attagacaaa taccacctgg gacattcagt catttgaaaa atcttgacac
301 tttaatgttg agtgataatt taatacaatc attagaaaat ggattgttta caggattgga
361 acagctacgc catctgtatc tttataaaaa caagattcgc aacattgaag agggagcgtt
421 tgatgggatg ccaaaattgg aacaacttta tttacactac aacaagataa tgaagataca
481 tccaaatttg tttaataacc tgaaaaattt acaaagactg tttcttcatg aaaacagaat
541 acattacatt cccagaggta ccttcgcaaa tatggcatat ttaaagagag tgcgtttgga
601 cggtaacgct ctagtatgtg actgcggact taaggaaatg atgatggcca tgccgaaggc
661 agaggtcgct gcagtctgtc agtatccgga gaacatgagg ggaaagactt tacacgaaat
721 gacacccagc gatatagtgt gcaacgaacc tcggattata gaaggaccac atgacgtgga
781 agtgagtttc ggcggttcgg ccattttcac ctgcagagtg gaagccgatt cggacgctga
841 aatcatttgg atgctcaaca gtaatgaaat tgacaaatcg gacccacgct actcggtcag
901 tcaggacggc acgttagtcg tgaagtctat gacggacaaa gaaatgggag tctatgaatg
961 tatggtgaag tcgacaaatg gaatgtccaa gtcacaggcc gctaagacct tatataatca
1021 tgtatcaggt aaagctgaat ggaaacgaaa gccgaaagat cagagtgcaa gagaaggtga
1081 caccgtacag ctagagtgtg ctgtctctaa cggacatgtg acttggactc gtaacggtgt
1141 accgattagt gcaccggtat cagcagaagg aacactgact ctatcttccg ttacacctca
1201 ggatacggcc agttacacat gtaacgtcga aacacctcag ggaaagatta aggccgaagc
1261 taagctaacg gtttcggcac ctccgaggat tttacaacca cctgagagtt catctgtaag
1321 aagtggagaa acggttttac ttcgctgtag agcaagtggt acgccagagc cgatattatc
1381 ttggtttaaa gatggtgcta gtctacaacc ttctgagaga ataatataca acgatgactt
1441 ttcagaagtc acgttgaaaa aagtttcttt aggagattca ggtttgtata cgtgtatggc
1501 tcagagtact atcggaacag ctgaagagag cggtgaatta agggtcaggc ctgtaggtcc
1561 cagacctcca gcgtttctta tcgctccgta tcctattata gcacaagcag gaacatcggt
1621 ggaactcccg tgtaaagtac agggtgatcc ttatccgaca atagaatggc tcaaaaatgg
1681 tgagccgatt cagctcgatc aaactcatag ggtatttcca atcggaagcc tccgccttta
1741 caatattaca caatcggatg cgggcatcta tagatgtaca gctacaaatg tgcacggtga
1801 aatatctgct cacgcaactc tcactgttga aggagcaaat aacatcgatg attcctttgt
1861 tcttacggca atacaaaatg ctaaagatag tgtcgatcga gccgttaatt caactttgga
1921 ctccatattt tcaaagaaag aacccaagaa cccgagtgca ttgatgcgac ttctcaggta
1981 tcccgatgaa caggcgagat cagtggtgcg agccgctgac atttatgaaa ccacattgtt
2041 aaacattaga aagcacataa taagtgggtt acaatttaat gaaacggaag attttcacta
2101 cgaagaggta ttgtctccgg accagttaca attaattgcc agaatgtccg gctgtaacga
2161 gcaccaaaaa gctaactgca gtaacatgtg tttccacact aaatatcgta cgatagacgg
2221 aacctgtaac aacttacaac acccgtactg ggcgcttca catacagcat ttagaaggtt
2281 gttaaaacct atttacgaaa atggatttag ccaacctgtt ggctgggaca aaaacagaaa
2341 atacaatggc ttcagccttc ctaatgctcg caaagtatca ttaacattaa taagaaccga
2401 aaacatcaca cacgacgata caataaccca tatggtgatg cagtggggac agtttctaga
2461 tcacgattta gaccatgcca ttccttcgac taccgccgag agctgggaag gtgtcgactg
2521 caagaagact tgtagagcta caccgccttg ttttccgatg gaggtcgaac aagacgatcc
2581 gagagtcaag aataggcgtt gtatggactt cattcgttct agtgccatat gcggttccgg
2641 aatgacctct gtgtttttca atacgattca accgagggaa caaattaacc agcttacttc
2701 cttcattgat gcgacacagg tttacggatt tcacgacaat agaagtatcc ttcttcgaga
2761 ttatacagaa gacttgggcc ttttgagagt cgggcttatt accgattcag gaaagcctct
2821 tcttccggta gctggtgcac aagaagtaga ctgcagaaga gatccgactg aaaacgaagc
2881 aggatgtttc ttagccggtg atatcagagt taatgaacag ataagtttgg tagctatgca
2941 cacaatttgg ctaagagaac ataacagaat agctacagag ttaaaagaaa tcaatccgca
3001 ttggaccgga gaaataattt tccaagaagc acgtaaaata gtcggagctc aaatgcaaca
3061 tattacctac aaacattgga ttcgaaatat tctcggagaa aaaggaatgc aaattctcgg
3121 agaatacgaa ggctatgatc ctaacttgga cgcaggaatt tcaaatgtat ttgcgacggc
3181 agcactaaga tttggacata ctctgataaa tccggtatta gcacgtctgg acagtgaatt
3241 taaaacgata ccagagggtg atttgccatt aggacgtgca ttttttgctc cttggagaat
3301 aatcgaagaa ggcggagtcg acgctttaat gagaggaatg tggatgtcgg cggctaaaaa
3361 gaaaatgcca aatcaaaatt taaataacga actgacagat catctcttta cgtcttttca
3421 cgctgtagcg ctagatctgg cttcgatgaa cgttcagagg tcgagagatc acggaatacc
3481 gttctacaat gagttcagga aattatgtaa tctatcttct gtccaaacgt ttgacgattt
3541 gaaagaacaa atatcaaatt ttgaagtgag agaaaagttg aaaaatcttt acggccatcc
3601 acacaacatc gacatatttg ttggcggaat actggaagac cagatcgacg gtgcaaaatt
3661 tggtcctaca ttccgctgtc ttctcatcga acaatttaag aggataagga acggtgacag
3721 attttggtat gaaaatcccg ctacattcaa agcgtcacag ttaacacaaa ttcgtcagag
3781 ttcactcgca agggttctat gtgacaacgg cgacaatata acaagaatat ctcccgatgt
3841 ttttatactc ccggaaaaac agacaccatc aattgtagac tgcagcgata taccaaaaat
3901 tgatctcaga ttttggtacg agtgcgaaga ttgtgaaaac gacgtatcat caagaggtag

```
     3961 acgtggtata tcaggcgaag atatggaaga gagaatggaa ggattagaaa acatggtaca
     4021 acaacttcag aaaaccgtaa aatccttaaa gaaaaaagta aacggtcttt caaagtgtcg
     4081 tgataataaa ggagttaaca ggaaggatgg agacacttgg caaaaggaca actgtacttt
     4141 gtgcgaatgt cagggtaagc aagtgtcttg tacaaggatt cagtgtgcca aactaacgtg
     4201 ccaaaggcaa gagaaagttg aagggagatg ctgcccggtt tgcgtttaat tattattatt
     4261 aatatttttg taaaatatat atttattact atgtactgtg tacagtgtcc aatgtaaaca
     4321 ctaagaatcg acgacttaat gtaaacattg gctacaacca atgtaaataa tataattaat
     4381 taataatgtg attcataatg cttgcttttt gtaataaaga ccgagagcta gatgttcagc
     4441 tagattataa gactttatta gtgttgtttt agcatatatt ttgtttgttt tttatttatt
     4501 gtgatatatt tctactatta ttttaattgt aatttgcagt ttttgttata attattgtat
     4561 taattaactt ttggg
//
```

```
LOCUS       Relish                  3478 bp    DNA     linear   UNA 03-MAR-2026
DEFINITION  Pyrrhocoris apterus Relish, complete mRNA.
ACCESSION
KEYWORDS    .
SOURCE      Pyrrhocoris apterus
  ORGANISM  Pyrrhocoris apterus
            cellular organisms; Eukaryota; Opisthokonta; Metazoa; Eumetazoa;
            Bilateria; Protostomia; Ecdysozoa; Panarthropoda; Arthropoda;
            Mandibulata; Pancrustacea; Hexapoda; Insecta; Dicondylia;
            Pterygota; Neoptera; Paraneoptera; Hemiptera; Prosorrhyncha;
            Heteroptera; Euheteroptera; Neoheteroptera; Panheteroptera;
            Pentatomomorpha; Pyrrhocoroidea; Pyrrhocoridae; Pyrrhocoris.
FEATURES             Location/Qualifiers
     5'UTR           1..158
                     /gene="Rel"
     gene            1..3478
                     /gene="Rel"
     mRNA            join(1..180,181..252,253..389,390..556,557..674,675..804,
                     805..909,910..1021,1022..1158,1159..1202,1203..1251,
                     1252..1319,1320..1398,1399..1497,1498..1658,1659..1851,
                     1852..2021,2022..2123,2124..2190,2191..2221,2222..3478)
                     /gene="Rel"
                     /note="Relish"
     source          1..3478
                     /mol_type="mRNA"
                     /organism="Pyrrhocoris apterus"
                     /db_xref="taxon:37000"
     CDS             join(159..180,181..252,253..389,390..556,557..674,
                     675..804,805..909,910..1021,1022..1158,1159..1202,
                     1203..1251,1252..1319,1320..1398,1399..1497,1498..1658,
                     1659..1851,1852..2021,2022..2123,2124..2190,2191..2221,
                     2222..2231)
                     /gene="Rel"
                     /product="Relish"
                     /translation="MSDIFYSDESGGYFSDIQYSPEPNANIHGNKEPFIEITEQPGDF
                     RYRYEKEKYHGHILGFTSPPTKKKDQCKFPAVKIHNYDKDVIIRCWLITTDLDEPKAH
                     FHKLARKPPKGTLMYEPHDLRATVRNNYEVTFDYLHIVHAVVSDENDLNIYKKKKNWI
                     RSRTYKNPSYTYEQGVDKEELKQMDKNQSILFFQAFDAVTEEKICEAVSRKIMNLHNP
                     ETAALKIVNYSRTYGTCKGRDEVFIFVEKISKDVEVRFFEINQATAQRSWQARADYGP
                     QHVHHQYGIVFRTPPFCDSNLSKDVLVYFELYRKTDGAKSEPVKYTYKPDAHCRMNND
                     FYKKRPHSPERFLNDTNNVLFVQPEIKRLREGRLSPDTTTLENNFGFPVEYNSDILNN
                     LTDQVSQEQYSWDTESQYHTLVTSDLDKYTQYNITSELQLIATTLPDFDSLLDGDNFC
                     TDSVPSSILKQPSEDILNLSSDEVSSSNEIPETVESEDIYVKTIKDFTEIRNIRDVIE
                     NFSDIKEAKKLLMSIAHINAVDFDGNSVLLVASECSLYALDVLLKHSEVDRSVTNLVG
                     DNALLCATRSKEEQLAEMLLQNKFDPNIQNYKTGETPLHVALSQENIGLVKLLLKYGA
                     DPLLNNYGSFSSYDYSMNCNSEELRRVIFRASQRNFKHRQTINIPEPPVNPMESELIE
                     ELESLNVS"
                     /codon_start=1
     primer_bind     1690..1709
                     /standard_name="Rel-Rev"
     primer_bind     complement(1769..1786)
                     /standard_name="Rel-Rev"
     3'UTR           2232..3478
                     /gene="Rel"
ORIGIN
        1 gtacttcaca ttatctgttt ctggttgcat cttctaccta tgtttctgat ttcagttcat
       61 attagtgtct gacctctgca cttgtcttat atgtttaagc tctaattttt atatttataa
      121 actcttcaaa ctttcaagga caatacattg tcgggaaaat gagtgacata ttctatagtg
      181 atgaatcggg aggatatttc tccgatattc aatattcgcc tgagccgaat gcaaatatac
      241 acggaaacaa agaaccgttt attgaaatta ctgaacaacc tggtgatttc cggtatagat
      301 atgagaaaga aaaataccat gggcacatcc ttggtttcac gtctccccct actaaaaaga
      361 aggatcaatg taaatttcct gcggtcaaga tacataacta cgacaaggat gttataatcc
      421 ggtgttggtt aataactacg gatctagacg agccgaaggc tcattttcat aaactagctc
```

481 ggaagccacc aaaagggact ttgatgtatg aacctcacga tcttcgtgct actgtgagga
541 ataactatga agtaactttc gattatctgc atattgttca tgctgttgtc agtgatgaaa
601 acgacttgaa tatctataaa aagaagaaaa attggatccg tagtagaaca tacaaaaatc
661 ccagttatac atatgagcaa ggagtagaca aagaagagct taaacaaatg gataaaaacc
721 agtccatttt gttttttcaa gcctttgacg ctgtcacaga ggaaaaaata tgtgaagccg
781 tttcaagaaa aataatgaat ttgcacaatc ctgaaactgc agcattaaaa attgtaaatt
841 acagtcgtac atatgggacc tgtaaaggaa gggacgaggt gttcatattc gttgaaaaaa
901 tttccaaaga tgtggaagta cgattttttg aaattaacca agcaacggcc caaagaagtt
961 ggcaagcacg tgccgactat ggcccacagc atgttcatca tcaatatgga attgttttca
1021 ggacacctcc attttgtgac tcaaatttat caaaagacgt actagtttat ttcgaattgt
1081 atcgaaagac agatggcgca aaaagtgaac ctgtgaaata cacctacaag cctgatgcgc
1141 actgtagaat gaataatgat ttctacaaaa aacgacctca ttcgccagaa agatttctta
1201 atgacaccaa taatgtctta ttcgttcaac ctgaaattaa aagattgcga gagggaagac
1261 tatctccaga tactacaact ctcgaaaata attttgggtt ccccgtggaa tataattcag
1321 atatcctaaa taatctaact gatcaggtgt cacaggagca atattcatgg gacactgaat
1381 ctcagtatca taccttagtg acttctgacc tcgacaaata cactcagtac aacattacca
1441 gtgagcttca gttgattgct actactcttc cagattttga ctctctgttg gatggtgata
1501 acttttgtac ggattctgtc ccatcttcga ttttgaagca gccttcagaa gatattctaa
1561 acctatcatc tgacgaagta tcaagttcaa atgaaattcc tgaaactgtt gaatctgaag
1621 atatttacgt taaaacaata aaagacttta ccgaaattag gaacattaga gatgtaatcg
1681 aaaacttttc ggacatcaaa gaagccaaaa agctgttgat gtccattgca cacattaatg
1741 cggtagattt tgacgggaat tcagtacttc ttgtcgcttc cgagtgctct ttatatgcac
1801 ttgatgtact tttaaagcat agtgaagtcg atagaagtgt caccaacctg gttggagata
1861 acgcactttt atgtgctaca cgtagcaagg aagaacagtt agcggaaatg cttttacaaa
1921 acaaatttga tccaaacatt cagaactata aaaccggaga aacacctctc catgttgcac
1981 tgagtcaaga aaatatcggc ttagtaaaac tgcttctgaa gtatggagct gacccacttc
2041 tcaacaacta cggatcattc tcttcctacg attattctat gaattgtaat agtgaagagc
2101 tgagaagagt aatcttcaga gcgtctcaaa gaaattttaa gcataggcaa actattaata
2161 ttcctgagcc acccgtcaat ccaatggaat ctgaacttat tgaagaactg gaaagcttaa
2221 atgtttcctg aacatgtgct atgtttatac tacaacttgg gatgtgtggt cgcatgggga
2281 gtagtaacct tcacacacta aacactgtgt ctgtctcaaa caagcttcta tcgatctcga
2341 cagagtttcg gtgtttggga ctctgcctat ttcctacaag aactaagcaa atgatacggc
2401 cttactatgc ggacctatta ttaaccaact caaacattac taaacaaaat taactcaatc
2461 tataaaaacc acgacctttg gccataaaga ccgtctaatg ttgttcaatt ccgattaagt
2521 aagtgtcata atttttctgt actctgtcaa tcctactact cacgttgtat acacatattg
2581 taaatatatc aataaagaga tttattagtg aaataatatt atataagact atagtttttt
2641 tatttatttt tataaatagt ttcttaagag ttgtatatat ttattatgtt atttgtaaaa
2701 gttttaatta tattttatat caataatttc tgtgatgtaa tgttgaattg taccttgtat
2761 gtctattggt gctaaaaaac tatttttata tatgggtcgt tcagttgaca cactggtcaa
2821 gtccattacc tgaaattttc aggaatgttg tttttatcat ttttgttgta taaaattatt
2881 tctgtatctc cgaatagaat tcaaatattt aatacatgag gagactcata tttttgtaag
2941 taaaatacat ctctcttaag tttatgttgt tcacacaact gaaagaggta atttattttt
3001 aataaaaatt ggactgccaa ctgtacggct caaataatat ttattttgag agaaaaattt
3061 gacctcgtaa atctttgatc atttttttgta catggtgggt attacacatt tcattttgta
3121 caaaatacca ggttcttaaa ataatggcag tttatttaga tttgtatatt aatgttggta
3181 cctgggattt taacatttgt attatttaag ttcaaattat aaaagtgcta caaattatgt
3241 attaccaaat ataattttaa tattaagtta acataatttt ttgaaagttt ttattaataa
3301 gtgaccatca atattgcatt ttttcacttc cctgatacct atttatgaaa tatatagtag
3361 gggaagtatt attttgtagt tatgctttga aaagaagatt atagttaaaa gatattcttt
3421 atctggcaaa aatcacccaa attgtccaaa atattgacga catttcaact tattttttt

//

```
LOCUS       Spatzle, isoform A        1885 bp    DNA     linear   UNA 03-MAR-2026
DEFINITION  Pyrrhocoris apterus Spatzle, isoform A, complete mRNA.
ACCESSION   urn.local...n-kbw23ki
VERSION     urn.local...n-kbw23ki
KEYWORDS    .
SOURCE      Pyrrhocoris apterus
  ORGANISM  Pyrrhocoris apterus
            cellular organisms; Eukaryota; Opisthokonta; Metazoa; Eumetazoa;
            Bilateria; Protostomia; Ecdysozoa; Panarthropoda; Arthropoda;
            Mandibulata; Pancrustacea; Hexapoda; Insecta; Dicondylia;
            Pterygota; Neoptera; Paraneoptera; Hemiptera; Prosorrhyncha;
            Heteroptera; Euheteroptera; Neoheteroptera; Panheteroptera;
            Pentatomomorpha; Pyrrhocoroidea; Pyrrhocoridae; Pyrrhocoris.
FEATURES             Location/Qualifiers
     5'UTR           1..31
                     /gene="spz"
     gene            1..1885
                     /gene="spz"
     mRNA            join(1..82,83..177,178..218,219..357,358..490,491..594,
                     595..1885)
                     /gene="spz"
                     /note="spatzle, isoform A"
     source          1..1885
                     /mol_type="mRNA"
                     /organism="Pyrrhocoris apterus"
                     /db_xref="taxon:37000"
     CDS             join(32..82,83..177,178..218,219..357,358..490,491..594,
                     595..850)
                     /product="spatzle, isoform A"
                     /gene="spz"
                     /codon_start=1
                     /translation="MAALESLSCVFATLLLFVDDGYSTPVGNTTSFPHREQKAGVKFP
                     LENKSTEWSRISHMNASRTPASGSDTPFVFPDDLPPIQFTPKVIKRSPACSEDNTFCE
                     SVDDYPEDYLQKMLSNKIDEYQAIMGVDELRVPDITQRIDGVDENPLCASVEQIVYPK
                     AAKNKDDKWLYVVNQPPYSQGVRIEKCLKTASSGSTSCLFIDQLPLGYKTFCKQKFIY
                     RKLVALDESGSTITDTFELPSCCSCTVQTNPLHSRLGLMKNPAKVTTQKPKTDS"
     primer_bind     612..631
                     /standard_name="spz-Fwd"
     primer_bind     complement(820..840)
                     /standard_name="spz-Rev"
     3'UTR           851..1885
                     /gene="spz"
ORIGIN
        1 agtcaagtga gaggatccaa gcccgaaggt tatggcagct ttagagagcc tctcttgcgt
       61 tttcgcaact ttactacttt tcgttgacga tgggtacagc acacctgttg gtaatacaac
      121 aagctttcct cacagagagc agaaggctgg cgttaaattt ccattagaaa acaaaagtac
      181 ggaatggagt aggatatcac acatgaatgc aagcagaact cctgcttcag gcagtgacac
      241 acccttcgta tttccagatg atcttcctcc tattcaattt actccaaaag taataaaaag
      301 atctccggct tgctccgaag ataataccttt ttgtgaatcg gtcgatgatt acccagagga
      361 ttacttacaa aaaatgttat cgaacaaaat tgatgaatat caagcgatta tgggtgtaga
      421 tgagcttaga gtacctgaca ttactcagag aatagatggc gtggatgaaa atccattatg
      481 cgcatcagtt gaacaaattg tttatcctaa agcagctaaa aataaagatg acaaatggct
      541 ttatgttgtc aatcagccac cctactctca aggagtgagg atagaaaaat gcttgaagac
      601 ggctagctct ggctccacct cgtgtctctt tatagaccaa ttgcctttgg gctataaaac
      661 attctgtaaa caaaagttca tctacagaaa acttgtggcg ttagacgaaa gcggttcgac
      721 gatcaccgac acttttgagt taccatcatg ctgttcgtgt acagttcaaa caaatccctt
      781 acactcgagg ctggggctaa tgaagaatcc ggcaaaagtg acgacacaaa agccgaaaac
      841 agattcataa atcttgagtc aattttacgt gtatatgtat tttgaatttg tgtaaatagt
      901 attataaatg tttaaaataa actgaaatat tggggttatt catttaaatg gtttaaccga
      961 cttctttgag aaagaagtaa atataaataa ctgacaattt tatagttttg attactaatt
     1021 tgattagttt ttttatttat tttttaaaga tgttgtgatt tttgtataaa tactttttta
     1081 ttttataatg atttcttgat tattttgtat gcctgtgaca tattgaagat tattttccta
     1141 ctttggctat atgtatatat acccatatat atatatatat atatatatat atatatatat
```

1201 atatatatat atatatatat atatatatat ccatatttat gatttttgta tattacaata
1261 acattgtgga atatgtggaa taaagaaaca taaattgttt ccgataagac ctctaaatct
1321 agtaaataat atgctaaggt agttttattt actaaacatt ctgtcaaacg tttcgcccac
1381 ggagggcatt ctcaagaaca ctgaaataca taattgtagg acatctgaaa ttttttttac
1441 aaaatttgtt tttattgcct gattatatta ttaccttgtt cttattatta tgtaccttag
1501 tatattatta aatgacattg tagataatca tttgtaataa tttaattaaa tcaatatgtg
1561 atgaatgctt atatgtgcac aaaatgatac agtaataata ctcaataatc tgacaatata
1621 tatatatata tatatatata tatacttgct aagtgctaag tcttatttaa gaaaaaatta
1681 tttaaacgaa gaaaattata attgttctgt gttatacata ctatattaat aataaaataa
1741 aacaaatgtg tctaatttat aagaacatca gtataatact ttgtagatgt atatttattt
1801 caatgtggca tgtaaacata tttgtattta aataattatg ctaccattgt tattttgaaa
1861 agaaaaaaaa agactgtact ttcta

//

```
LOCUS       Suppressor of cytokine signaling at 36E        3557 bp    DNA     linear
UNA 03-MAR-2026
DEFINITION  Pyrrhocoris apterus Suppressor of cytokine signaling at 36E,
            complete mRNA.
ACCESSION
KEYWORDS    .
SOURCE      Pyrrhocoris apterus
  ORGANISM  Pyrrhocoris apterus
            cellular organisms; Eukaryota; Opisthokonta; Metazoa; Eumetazoa;
            Bilateria; Protostomia; Ecdysozoa; Panarthropoda; Arthropoda;
            Mandibulata; Pancrustacea; Hexapoda; Insecta; Dicondylia;
            Pterygota; Neoptera; Paraneoptera; Hemiptera; Prosorrhyncha;
            Heteroptera; Euheteroptera; Neoheteroptera; Panheteroptera;
            Pentatomomorpha; Pyrrhocoroidea; Pyrrhocoridae; Pyrrhocoris.
FEATURES             Location/Qualifiers
     5'UTR           1..359
                     /gene="Socs63E"
     gene            1..3557
                     /gene="Socs36E"
     mRNA            1..3557
                     /gene="Socs36E"
                     /note="Suppressor of cytokine signaling at 36E"
     source          1..3557
                     /mol_type="mRNA"
                     /organism="Pyrrhocoris apterus"
                     /db_xref="taxon:37000"
     CDS             360..1721
                     /product="Suppressor of cytokine signaling at 36E"
                     /gene="Socs36E"
                     /translation="MGQKFSDLKGIFGIMDEKDVNEVHISVESVDVCLRNNDESSAIC
                     RSSDDNGNSEPPSSINEHFDRLYITSEIREQEENGNIVPEASCSSQQVCTSTQSLFEN
                     KKSKRHKRGKNKEPPTTSGKKKRSHWVLRFNCARLKSGSNSSGASSEPDIAEPNIGLE
                     SCVCTGYRRTEDHHLGAGVVFEASSRRQYSGMSSESDDELHQIIGLDKFRADEYSLED
                     CDERARLERAREMAEGVDPPPGFRPAQHIQLVCPQDMNIDSLTALLHSAALNALSQLD
                     SQRVVHTQVDYIHCLVPDLLQITVCSFYWGKMDRYEAERLLDGKPEGTFLLRDSAQEE
                     YLFSVSFRKYGRSLHARIEQWKHLFSFDSHDPGVYSSPTVCGLIEHYKDPSCCMFFEP
                     MLTIPLHRNFAFPLQHLARAVICSKISYDGISQLKLPKPLKSYLKEYHYKQRVKVRRF
                     DTE"
                     /codon_start=1
     primer_bind     694..716
                     /standard_name="Socs36E-Fwd"
     primer_bind     complement(884..903)
                     /standard_name="Socs36E-Rev"
     3'UTR           1722..3557
                     /gene="Socs63E"
ORIGIN
        1 agtgaatctg acttgctttc caaaatggtg tcaataatgt tatttgtgtg attatttatt
       61 tttttatttt catcttttgt ttgtgttgaa agtgttatgt ttcttacaat gacttatttt
      121 agttgtggta atttaaatat gaattattct ggaagatgat ttattggtgc aatacgttgt
      181 ccttaaatag aaattgtcag acggacttca ctgcgctgca aaaaagtgat taccttcttt
      241 gtttttttctc ttatttagtt ttatttactt gtacttctcc acatcgtatt aataatggtt
      301 gatcaatgga ttattggcct ttcagaattg ggcctttgga agtttttcat tgaaagataa
      361 tgggacaaaa gtttagtgac ttaaagggaa tttttggtat aatggatgaa aaagatgtca
      421 atgaagtgca tatttcagtg gaaagtgtag atgtctgcct aagaaataat gatgaaagtt
      481 ctgcaatttg cagaagtagt gatgataatg ggaatagtga accacctagc agtataaatg
      541 aacactttga taggctttat attacttctg aaattagaga acaagaagaa aacggaaata
      601 ttgttcctga agctagttgt tcttctcaac aagtgtgtac atccactcaa agtttgtttg
      661 aaaataaaaa gtctaaaagg cacaaaagag gaaagaataa agaacctccc accactagtg
      721 gaaagaagaa acgttctcac tgggttttga ggtttaattg tgctcgttta aaaagtggta
      781 gcaacagcag tggtgctagt tcagagccgg atatcgcaga acctaatatc ggattggagt
      841 catgtgtctg tactggttat agaagaacag aggaccatca tctaggtgct ggtgtagttt
      901 tcgaagcatc atctcgccgg cagtattcag gaatgagcag tgaatcagat gacgagttgc
      961 accagatcat tggactggac aaatttagag ctgatgaata ttctctggaa gactgcgacg
```

```
1021 aacgtgctcg actagaaaga gctagagaga tggcagaagg ggtcgaccca cctcccggct
1081 tcagaccagc tcaacatatt caacttgttt gtccccagga tatgaatatt gacagtttga
1141 ctgcactcct ccattctgct gctctcaatg cactttctca gttggattcc caaagagtgg
1201 tgcatactca agtcgattat attcattgtc ttgtaccaga tcttttacaa attaccgtgt
1261 gttcattcta ttggggaaag atggatagat atgaagcaga gagactcctt gatggaaaac
1321 cagaaggtac atttttactt agagattccg cgcaagaaga atatttattt tcagttagct
1381 ttagaaaata cggaaggtcc cttcatgcta gaattgagca atggaaacac ttgttcagtt
1441 ttgactcgca tgaccctggt gtttattctt ctccgactgt gtgtggcctt attgaacatt
1501 ataaagatcc gtcgtgctgt atgtttttcg aaccgatgct tactatacct cttcacagga
1561 attttgcttt cccacttcag catttggcta gagctgtcat atgtagcaag atttcatatg
1621 acggcataag ccaactgaaa cttccgaaac cattgaagag ttatttgaag gaataccatt
1681 ataaacagcg cgtgaaagtt cgtaggtttg atacagaata attaattgag aaatcatcac
1741 taaatacatt ttaaagccaa atggtgcatt acatttagac aacttacaat cagatatcga
1801 ggtttatatt tgttatatat attattaatt actccttatt ttgattgcct gctgtcctat
1861 ttaaactgtt aaatatatac gatgttcctg ttattttaat ggtaatgatt tggaactgtt
1921 tattatgtat ttagaagaat actcaaagaa acaacagtat tagctattga agtgattata
1981 tgatattcac ttttattgta tgggtgtcta ttttagatat tatttttttg aacttgaatt
2041 tcatttttaa aatgtaaatg aaaagttaaa attccaaaaa ttaaaaaaaa atatatctaa
2101 agtgtttgtt atatgaggtt tgaatattta aataaaaagt ttatataaaa caaaaaaaaa
2161 tgtttattgt ataattgaaa aattttgcat ttagttaata tataattcat ttatataaat
2221 ttatggacac agataactta ttgttgtttt tttttgtctt atcagctatt acttattaat
2281 acttttgaaa tagttgcaag aggattaatg tgtttgatat aatctgcttt gaagttatta
2341 ttaattgcga agtatataca gttataatgt attatttcac agttttcact gttatcatta
2401 ttatgttgta atatttctta atattatcat tttttcagtc aggattatta aaaaagtgat
2461 tagttgttaa aatctatgtc atgcccatat tttaccttaa catgtaggta aggaatgggt
2521 gtagcaaaat gtcaatgctt ttttaaaaca gaatcatgtt gtcttttgat tgtttttgaa
2581 tcattgttta gaaatttgat acaaacaata accgaaaatt ctaataaaaa tgttttagta
2641 tggcttgttt aaatttatgg tcggaatatt ttcttccaat tgtattattt tgttacgaaa
2701 gaaatatttg tttttttaaa acactttttg tgtttaaaca ttaatatcta aacttaagcc
2761 gggcgtaata tttttttctt ttgttaatag aagtagtaaa agttgtcatt tgttatgaaa
2821 tttttcagta tctctcacat ttaatatata tgaagacaga atttcttttt aaattattat
2881 taatttgaaa ggacgtgtaa ctcagtagct gagttttatt taacgtatta cctaagcata
2941 tctataggat ttcgttttcc aaaagtataa aacaagtcca tatttgatta tggcttttga
3001 ttattaaatt tgatacttgt aaatgttcct taaagaatca agtatctgtg tataattata
3061 atagattgta tatctatata tacactagta cttttgcagc tttacgaata aaccttgtgt
3121 ggctttagat ttatatagta atatattcac tcgtaataat acatgttagt gtggttaaga
3181 ctgattcacc gttaaaaacg tttgtaatac cgaacaccat ttttaaaata tattgagaat
3241 tgctggtata atctcgtaat atgtgataat attgtaagtt caagtttcat ttaaataaac
3301 aaattgaatt aaaataattc ataacaagag ttaaatcgaa tcatttatag tatttattat
3361 ctaatcataa tacgttcctg tttttattaa caggatgata atgccatcat caattgtggg
3421 atacagacaa ataatattta catttgaaat tcttaatctt aagtaacaaa cattaattta
3481 ttttattggt tgtcattgtt tatttgttat acagttacaa atataattct aaattaactc
3541 cgattgtctt aatttaa
//
```

```
LOCUS       Toll, isoform A        4049 bp    DNA     linear   UNA 03-MAR-2026
DEFINITION  Pyrrhocoris apterus Toll, isoform A, complete mRNA.
ACCESSION   
KEYWORDS    .
SOURCE      Pyrrhocoris apterus
  ORGANISM  Pyrrhocoris apterus
            cellular organisms; Eukaryota; Opisthokonta; Metazoa; Eumetazoa;
            Bilateria; Protostomia; Ecdysozoa; Panarthropoda; Arthropoda;
            Mandibulata; Pancrustacea; Hexapoda; Insecta; Dicondylia;
            Pterygota; Neoptera; Paraneoptera; Hemiptera; Prosorrhyncha;
            Heteroptera; Euheteroptera; Neoheteroptera; Panheteroptera;
            Pentatomomorpha; Pyrrhocoroidea; Pyrrhocoridae; Pyrrhocoris.
FEATURES             Location/Qualifiers
     5'UTR           1..274
                     /gene="Toll"
     gene            1..4049
                     /gene="Toll"
     mRNA            join(1..273,274..1455,1456..1759,1760..2386,2387..2551,
                     2552..4049)
                     /gene="Toll"
                     /note="Toll, isoform A"
     source          1..4049
                     /mol_type="mRNA"
                     /organism="Pyrrhocoris apterus"
                     /db_xref="taxon:37000"
     CDS             join(275..1455,1456..1759,1760..2386,2387..2551,
                     2552..3385)
                     /gene="Toll"
                     /product="Toll, isoform A"
                     /translation="MKLLLLLLLHLIPLVLSKVQCPGSANCSCSNGVSGDFEILCPEN
                     ISAQPTIVTTFQPKNYIRIQCSKANSWKELELMSGLNLGPVKFFVLMLCPLPGISFNE
                     LMTTMGIPSIKSLQFQSFGNISNTLTKEHLEGLHDLNTLIFYNNDLTELPEDLFKDVG
                     NLISLLLNYNQVVLPKGIFRHVPKLEVLELGSNNISFLEPGIFRNLTKLRLLNLWGNR
                     LQNLTRSVFSDLQNLEGLDLNNNGLKTLPPDVFTDLIKLKNINLYNNDFVSLPQSLFR
                     SSSRMEIIQINSNRHTLKTLPPAFFANLTNLKYLNLKNNNISELPEDLLIGCMALDLV
                     ELQYNNLRELPEKIFHDIQNATRIDLSYNKLVNLPNDLLSSLNKLKTLNLSHNEITQI
                     QEDLFSNLYELQIIKLSHNKIRTIHSEAFKATTHLHTVDFSYNELIELGNSLIFNASP
                     LKLATNLEYLNLSHNNLSYFYNDWQISMTKLQFLDMSYNNFTKLMIHDIQQFTSTSLT
                     VDMSHNNITEVSLDEAESLAAELPILHGVRENNSPKVILRANPLQCDCKAYELIRYFR
                     QHLEPEIYLLATFDGTNLACAGPEELKDTLVMNLDPREVACKANEMECPKNCSCYYRP
                     SNSAMIVDCSRRNLTRIPPVMPTNGTLFSKTNHTELDLKGNNLVTFPDRLGPGYSNVT
                     KLYLSYNNLTSINITSFSKDLQVIELDNNNLTRMDTGSLKTLEKLGHLEIITLQNNPW
                     ICDCQSKEFHSFLQRNYKKVPLLENITCSNGALVTKLTVYDLCPVTHILIKAASLVVA
                     VIGLIIGASIGFYYRYQKEIKVWLYAHRCCLWFVTEEELDKDKKYDAFVSYSHQDEKF
                     IIEHLQPVLEKGDPKYSLCLHYRDWLVGDLIPDQIARSVEDSRRTIVILSPHFLESVW
                     GKMEFRTAHCQALREGRARVIMILYGDIGPTDKLDPELKAYLSTNTYVKWGDPWFWQK
                     LRYALPHPAKHSKGTAVQLIPRQSILQNGTEKLINGSDPNSTSVTTPPANALIIEPFK
                     ADSKPI"
                     /codon_start=1
     primer_bind     3002..3021
                     /standard_name="Toll-Fwd"
     primer_bind     complement(3210..3228)
                     /standard_name="Toll-Rev"
     3'UTR           3386..4049
                     /gene="Toll"
ORIGIN
        1 agtcctgtgc cactcgaggt acgctaaact attctcgcgt tcagaaacaa aatacatcgc
       61 ccttttttaaa tgtttaatta aataaaaata tttatattac atctaatgtt caaaaagtgt
      121 tgtagtcata gaaaataata atacaacaaa tattattttt ttgtatttca agtgcttagt
      181 aaattaaata aatcaactgt aattaaaaca ttcaaccaga tgacaacata gtgaagacat
      241 catcaccatc ggttgacaag acaacatatc aaggatgaag ttattattat tactcctact
      301 gcatctaatt ccattggtgc tgagtaaggt ccaatgtcca ggaagcgcta actgttcctg
      361 ctcgaatgga gtgagcgggg actttgaaat tctctgccca gaaaacattt cagctcaacc
```

```
 421 tactatagta acaacatttc aaccaaagaa ttacattagg atacaatgtt caaaggcaaa
 481 cagctggaaa gaattggagt tgatgagcgg ccttaacttg ggtcctgtaa agttttttgt
 541 tcttatgctc tgccctctac ctggtatatc gtttaacgaa ctcatgacta ctatgggaat
 601 accttccatc aaaagtttac agtttcaatc gtttggcaac ataagcaata cactgacaaa
 661 agaacactta gaggggcttc acgacttaaa tacgcttatc ttctataata atgacttaac
 721 agagctaccg gaagatcttt ttaaagatgt tggaaacctt ataagcctac tgcttaatta
 781 taatcaagtt gtattaccaa aaggaatttt tagacatgta ccgaagcttg aagttctgga
 841 attaggaagc aataatatta gcttcttgga gcccggtatt ttcagaaatt tgacaaaatt
 901 aaggctttta aatctttggg gaaatagact tcaaaacttg acgagatcag tttttctgta
 961 cttgcaaaat ttggaaggtt tagatttgaa caataatgga cttaaaacac ttccaccaga
1021 tgtttttacc gatttgataa aactaaaaaa tatcaacttg tataacaatg attttgtttc
1081 gttgccacaa agcttgttcc gcagttcgtc ccgaatggaa ataatacaga ttaactcgaa
1141 tcgtcataca ttaaaaacat tgcctcctgc attttttgca aacttaacga atttgaaata
1201 tctaaatttg aaaaataata acatatctga gcttccagaa gatctactaa taggctgtat
1261 ggctcttgat ttggtagaat tgcagtacaa caacctacga gagctcccag agaaaatctt
1321 ccacgacatt caaaatgcca caagaataga tttatcatat aataaactgg ttaatctgcc
1381 gaatgatttg ctctcatcat taaataagct aaaaactctt aatctgagcc ataatgaaat
1441 cactcaaatt caagaggatt tattttcaaa tttatatgag ctgcaaatta ttaaactttc
1501 tcacaacaaa attaggacaa ttcattcaga agcctttaaa gctacgacac atttacacac
1561 tgtagacttc tcgtataatg aactcattga attaggaaat agtttaatct ttaatgcctc
1621 tcctctaaaa ttagcaacaa acttggaata cttaaactta tctcacaaca atttaagtta
1681 tttttacaac gattggcaaa tttcaatgac aaaacttcaa tttctcgata tgagttacaa
1741 caatttcaca aagctaatga ttcacgatat acaacaattt acatcaactt ctctgacagt
1801 cgatatgagc cataacaata tcacagaggt atcactcgat gaagccgaaa gtcttgcggc
1861 cgaactacca atacttcatg gagtccggga gaataacagt cccaaggtta tattacgagc
1921 gaatccgtta cagtgtgact gtaaagcgta cgaattaatt cggtacttca gacaacattt
1981 ggaaccggaa atatatcttc ttgcaacatt tgatgggaca aatctagcct gtgccggccc
2041 tgaagagtta aaagacacat tagttatgaa cctcgatccg agagaagttg cttgcaaagc
2101 aaatgaaatg gaatgcccta aaaactgctc ttgttattat agaccgagta attcagcgat
2161 gattgtcgat tgttctcgaa gaaatctcac aagaatacca ccagtgatgc caacaaacgg
2221 aacacttttt tcaaaaacca accacaccga attagatctt aaaggaaata acctggtaac
2281 tttccctgat cgacttggtc cgggctattc caatgttaca aaactgtatt tgtcctacaa
2341 caatctgacc agcattaaca ttacaagctt cagcaaagat ttacaggtta tagaacttga
2401 taataacaat ttaacacgaa tggacaccgg atcactgaaa acactagaaa aattaggaca
2461 tcttgaaata atcactttgc agaataaccc ttggatatgt gactgccagt ctaaggaatt
2521 tcattcattt cttcaaagga attataaaaa ggtgcctttg ttagaaaaca tcacgtgctc
2581 aaacggtgct ttagtgacta agctgacagt atatgatctg tgtcctgtaa cacatatact
2641 aataaaagca gcgagtcttg tggtagcagt aataggactc atcataggag catcaattgg
2701 attttactat cgataccaga aagaaataaa ggtttggctg tacgctcacc gatgctgcct
2761 gtggtttgta acagaagaag agcttgataa ggataagaaa tacgacgctt tcgtaagtta
2821 ctctcatcag gatgaaaaat ttattatcga acatttacag cctgtcctcg aaaaaggcga
2881 tccgaagtat tcgctttgcc ttcattatag ggactggctc gttggtgatc ttataccgga
2941 tcagatagca agatcagtcg aggattcgcg aaggactata gttatattgt cgccacactt
3001 tttggaaagt gtctggggta aaatggaatt caggaccgcc cactgtcaag ctctacgcga
3061 agggcgggct agagttataa tgatacttta cggtgacatt ggaccgacgg ataagctgga
3121 tccagagtta aaggcttacc tttcgacaaa tacgtacgtt aagtggggcg atccgtggtt
3181 ttggcagaag ttgcgctatg cccttccgca cccagccaaa cattccaagg gaacagcagt
3241 tcaacttatt ccgagacagt caatcttaca gaatggtact gaaaagctta taaacggatc
3301 cgatcctaac tcgacttcgg taacaacacc tcctgcaaat gcccttatca tcgagccttt
3361 caaagcggac tctaagccca tataattatg tatacacaca aattgaccag tttgtctaaa
3421 gtgttttagc agtgaaagtg actagatatt ttaactaact gatacaattt gtggtaatta
3481 aaagtatctt ttcgaggaac taagcttttg tgtatgatga aaagactaat tatggttaaa
3541 ttataatgta tttgtgagta tatgtgttct ttaaatattt taaagaagtg tttgatgtgt
3601 gtataaacaa tataaatgaa aaccataatg attagttcga tactaattgt tgtacgccta
3661 atttataatg tacaaaattt tttagattga tactcatata tatattactg tgatacaaat
3721 ctcatttaaa taaatatttt tacctaatta aaagactaca cataaattta tgtgattcaa
3781 taatgtacaa ttgtacacaa gactgtatgt aaatttagaa ttaatgtaaa tagttaagaa
3841 tgttagctca atgtctagta atttaatttt taagtattga attgtttgta tttgtcagtt
3901 ttaagtgcca tgattgtttt gtcttgctta gaatgtagac ctattcagat caactttgtt
3961 gttgtttttt tttatctaag gaacaaaata aaaagtatgt aaatattaaa tgtcaataaa
4021 ttatacaaaa tttcctacct tttctaaaa
//
```
